# Spatiotemporal Cascade of *AP1*/*CCA1*-*TOC1* Module Gates Stem Development

**DOI:** 10.64898/2026.05.30.728956

**Authors:** Jiahui Xu, Jiahong Qi, Yufeng Yang, Chuan Jiang, Pedro Garcia-Caparros, Yichao Zheng, Mei Jin, Qingyu Guo, Dan Zhao, Lin Guo, Yunzhen Li, Xiaoli Fan, Yuehui He, Xiaodong Xu, Qiguang Xie, Xigang Liu, Hao Zhang

**Affiliations:** Hebei Research Center of the Basic Discipline of Cell Biology; Ministry of Education Key Laboratory of Molecular and Cellular Biology; Hebei Collaboration Innovation Center for Cell Signaling and Environmental Adaptation; Hebei Key Laboratory of Molecular and Cellular Biology; College of Life Sciences, Hebei Normal University, Shijiazhuang, Hebei 050024, China; Department of Superior School Engineering, University of Almería, Ctra. Sacramento s/n, La Cañada de San Urbano, 04120 Almería, Spain; College of Life Sciences, Hengshui University, Hengshui, Hebei 053000, China; National Key Laboratory of Crop Genetic Improvement, Hubei Hongshan Laboratory, Huazhong Agricultural University, Wuhan, Hubei 430070, China; Chengdu Institute of Biology, Chinese Academy of Sciences, Chengdu, Sichuan 610213, China; Peking-Tsinghua Center for Life Sciences & State Key Laboratory of Wheat Improvement, School of Advanced Agricultural Sciences, Peking University, Beijing, 100871, China; State Key Laboratory of Crop Stress Adaptation and Improvement, School of Life Sciences, Henan University, Kaifeng, Henan 475004, China

**Keywords:** Stem development, *AP1*, *CCA1*, *TOC1*, Pectin methylesterase, Spatiotemporal regulation, Plant height

## Abstract

Stem development is crucial in plant vertical architecture and overall crop improvement. The molecular mechanisms underlying the initiation and circadian growth of stem remain still enigmatic. Here, we demonstrate that *APETALA1* (*AP1*) spatially initiates circadian stem growth through a noncell-autonomous role of *TIMING OF CAB EXPRESSION1* (*TOC1*) emanating from the floral meristem to inflorescence meristem in *Arabidopsis thaliana*. Mechanistically, AP1 interacts with and recruits CIRCADIAN CLOCK-ASSOCIATED1 (CCA1) to associate *TOC1*. Dynamic formation of the AP1-CCA1 complex is crucial for maintaining the circadian rhythmicity and expression of *TOC1*. Tissue-specific expression of TOC1 in floral meristems rescues the developmental defects of *toc1* mutants at both functional and transcriptional levels. Furthermore, TOC1 directly activates the circadian expression of pectin methylesterase gene family, which is critical for its role in stem development. *AP1* homologs regulate circadian stem elongation and plant height in wheat and rice, underscoring the conserved mechanism across flowering plants. Our findings uncovered a spatiotemporal regulatory cascade gating stem development and identified candidate genes for crop improvement.

## INTRODUCTION

One of the most successfully evolutionary events for modern land plants is the development of a vertical shoot axis (stem), which enables plants initiating the transition from two-dimensional to three-dimensional growth to establish optimal shoot architecture^1–3^. Shoot architecture is mainly determined by the activity of meristems, including the shoot apical meristems (SAMs), and the subsequent development of various organs^4^. Orchestrated with organogenesis, stem development, including stem initiation and growth, is a key factor for shoot architecture formation, yet it has been less understood compared to the development of leaves and inflorescences^2,5^. During the vegetative phase of *Arabidopsis thaliana*, stem development is inhibited prior to the floral transition but becomes activated during this transition. Subsequently, stem growth undergoes in a rhythmic pattern, with active cell proliferation near the apex playing a dominant role in stem growth (Extended Data Movie 1)^2,6^. Despite its importance, stem development has been relatively neglected compared to other organs^2,7^. Meanwhile, coordinated stem development and organogenesis are crucial for crop improvement. Changes in phyllotaxis and stem height can significantly affect photosynthetic efficiency and grain yields^8–10^. For instance, impaired gibberellin (GA)-related dwarfing genes in wheat and rice led to semi-dwarf plant and consequent increasing grain yield during the Green Revolution^10–13^. Moving forward, alternative strategies to decrease plant height by leveraging other phytohormones like brassinosteroids and targeting key developmental genes involved in stem development may contribute to the next Green Revolution^14,15^. However, the underlying developmental mechanisms governing stem initiation and growth remains unclear^2^.

All aerial organs in plants are derived from stem cells located within the SAMs^16^. The SAM is structurally organized into several distinct zones: central zone (CZ) harboring stem cells, a surrounding peripheral zone (PZ), and the rib zone (RZ), located below the CZ, including the rib meristem (RM) and the subapical peripheral region that supports ongoing stem development^5,7,17^. The outermost cell layer (L1) and the adjacent cell layer (L2) undergo anticlinal cell divisions, ensuring the maintenance of the meristem organization, while cells of RM (L3) undergo transverse divisions to form the pith (Fig. 1a) ^5,7,17–19^. During the vegetative phase, the CZ and PZ sustain the formation of leaf primordia, while RM is inhibited. During the floral transition, the RM is activated initiating rapid stem elongation, in which the homeodomain protein REPLUMLESS (RPL) locally controls RM elongation^2,5,7^. Nevertheless, the precise mechanism by which the stem initiation occurs and the rhythmic stem growth is regulated remain largely unknown.

**Fig. 1.**
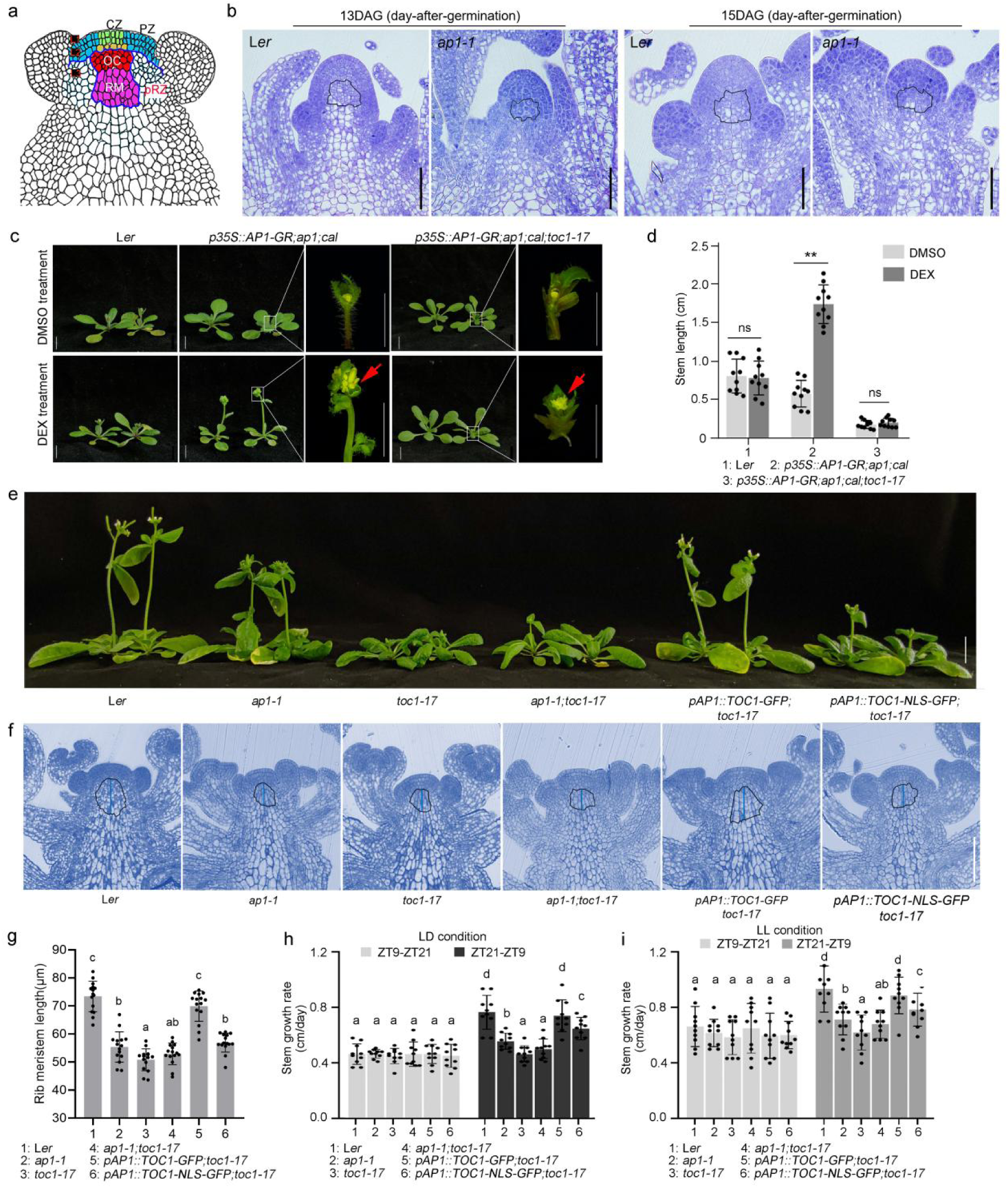
*AP1* and *TOC1* control stem initiation and circadian growth. (**a**) Schematic representation of the shoot apical meristem composed of the central zone (CZ), a surrounding peripheral zone (PZ), organizing center (OC), and the rib zone (RZ), including rib meristem (RM), and a peripheral rib zone (pRZ). (**b** and **f**) Longitudinal sections of shoot apex of indicated plants during floral transition (**b**) and reproductive stage (**f**) showing the length of Rib Meristem (RM) (**f**). The black line outlined the RM region and the blue line marked the RM length. Representative section from 15 samples per genotype was displayed. (**b**) Scale bars: 50 μm. (**f**) Scale bars: 100 μm. (**c**) Stem development of indicated plants treated with DMSO and DEX, respectively. Plants at 13 DAG (days after germination) were treated with 1 μM DMSO or DEX every two days for one week. Inserts show the shoot apexes with rosette leaf removed. Red arrows mark floral organs. Scale bars: 1 cm. (**d**) The stem length of indicated plants after treatment with DEX and DMSO, respectively. Data represent mean ± SD (n=10). Statistical significance was determined by Student’s *t*-test (two-tailed); ns denotes not significant; **p < 0.01. (**e**) Morphological observation of indicated plants. Scale bar: 1 cm. (**g**) Statistical analysis of RM length of indicated plants. Data represent mean ± SD (n=15). Different letters indicate statistically significant differences by one-way ANOVA (Tukey’s multiple comparisons test, P < 0.01). (**h** and **i**) The stem growth rate in night and day of indicated plants under different conditions of LD (12h light/12h dark) (**h**) and LL (constant light) (**i**). ZT, zeitgeber time. Data represent mean ± SD (n = 10). Different letters indicate statistically significant differences by one-way ANOVA (Tukey’s multiple comparisons test, P < 0.01).

The floral transition is regulated by multiple signaling pathways that are ultimately integrated by floral meristem (FM) identity genes: *LEAFY*, *APETALA1* (*AP1*) and *CAULIFLOWER* (*CAL*)^17^. *AP1* and *CAL* are homologous genes encoding MADS domain-containing transcription factor that are both necessary and sufficient for the transition from inflorescence meristem (IM) to FM^20^. The expression of *AP1* and *CAL* is spatially restricted to the FM primordium, and *ap1;cal* double mutants produce massively over-proliferated IM-like meristems^21,22^. During early flower development, *AP1* orchestrates a cascade of gene expression that promotes both floral initiation and floral organ patterning with either repressor or activator activity^22–25^. Despite these well-characterized roles in floral development, the potential involvement of *AP1* in regulating stem development remains enigmatic.

The cell wall matrices are the outmost layer of growing plant cell, which are constituted of multiple polysaccharide polymers including cellulose, hemicelluloses and pectins, as well as small amounts of structural proteins^26^. Pectins primarily consist of homogalacturonan (HG), rhamnogalacturonan I (RG-I) and RG-II^27^. HG is an unbranched homopolymer of α-1,4-linked D-galacturonic acid, initially synthesized in a highly methyl-esterified state and then de-esterified by pectin methyl-esterases (PMEs). This de-esterification is inhibited by pectin methyl-esterase inhibitors (PMEIs) in cell walls^28^. Demethyl-esterified HG triggers two distinct cell wall changes: it is either degraded by polygalacturonases (PGs) to induce wall loosening, or alternatively cross-linked by calcium ions to reinforce wall rigidity, in which PME processivity determines these differential outcomes. Distinct SAM domains differ markedly in growth dynamics, cell geometry and mechanical traits, determined primarily by cell wall composition and remodeling. Enzymatic modification of cell wall homogalacturonan is critical for SAM mechanical properties and organogenesis^29,30^. In the SAM, organ primordia have low levels of pectin methylesterification and highly extensible cell walls, whereas the stem cell domain exhibits high methylesterification and comparatively rigid walls^30,31^. Local PME activity drives lateral organ emergence, while widespread PME suppression restricts cell wall extensibility and blocks organ formation, demonstrating that pectin modification status is central to meristem maintenance^32^. While several studies revealed the roles of PMEs such as *PECTIN METHYLESTERASE5* (*PME5*) and *PME VANGUARD1* (*VGD1*) in cell wall patterning and consequent meristem maintenance and organ outgrowth^32–34^, the functions of PMEs in stem development still remain unclear.

To adapt to the 24-h d/night cycle, species have evolved endogenous circadian clocks that integrate cyclic environmental signals to achieve spatiotemporal synchronization through core oscillator genes^35–37^. Notably, circadian rhythms can vary significantly between individual organs and even among regions within the same tissue^36,38^. In plants, the circadian clock in the shoot apex acts analogously to the suprachiasmatic nucleus neurons in mammals, modulating circadian activity in distal organs such as roots and showing that the plant clock is organ-specific^39,40^. Recently, clock-controlled spatiotemporal florets development in sunflower was well studied^41^. Despite these insights, the mechanisms by which the circadian clock regulates SAM maintenance and activity remain largely unclear. Among the core circadian oscillator components, *CIRCADIAN CLOCK-ASSOCIATED1* (*CCA1*) and *TIMING OF CAB EXPRESSION1* (*TOC1*) are key components of the circadian oscillator, representing morning- and evening-phased genes, respectively^42,43^. CCA1 directly represses *TOC1* expression, and conversely, TOC1 represses *CCA1* expression to form a feedback regulation loop critical for the maintenance of the circadian clock function. CCA1 initiates and sets the phase of clock-controlled rhythms by directly binding to and regulating its target genes. Conversely, TOC1 generally functions as a transcriptional repressor, directly binding to and repressing approximately all key oscillator genes, while recent study revealed its activator activity^44–47^. Although it is well known that rhythmic hypocotyl elongation and growth are controlled by light signaling and circadian clock components such as CCA1 and TOC1^48–53^, the extent to which the *CCA1*-*TOC1* module regulate stem circadian growth remains unknown.

In this study, using the model plant *Arabidopsis thaliana*, we uncovered a spatiotemporal regulatory cascade that gates stem development. Specifically, AP1 interacts with CCA1 to maintain both the expression level and rhythmicity of *TOC1* within the FMs. This regulation acts non-cell-autonomously and spatially activates RMs and initiates stem development through regulation of TOC1 on the expression of PMEs.

## RESULTS

### *AP1* activates stem initiation and sustains circadian stem growth during floral transition

To understand the dynamics of stem development, we tracked the structural changes of SAM and stem during the floral transition in Landsberg *erecta* (L*er*) wild type (WT) plants. Usually, the lower boundary of the RZ is hard to discern due to the unclear anatomical cell characterization^2^. Based on longitudinal sections though the shoot apex, we outlined the RM region according to the first elongated cell and the daughter cells detached from each other (gaps between the cell vertices), in which the RM cells characterized by the small transversal-divided cell with dense cytoplasm (Fig. 1a,b). At 11 days-after-germination (DAG), the RM was quiescent and surrounded with few RZ cells resulting in inactive stem development (Extended Data Fig. 1a). During floral transition at 13 DAG, the CZ, PZ and OC of the SAM were stable, except of the emergence of FM primordia instead of leaf primordia. Meanwhile, RM cells were activated triggering cell division and growth in RZ to initiate stem development forward (Fig. 1b), demonstrating that the RM activity and correspondent stem development were activated during floral transition^2,7^.

We then hypothesized that noncell-autonomous signals from FMs spatially induce RM activation and promote stem growth during floral transition to orchestrate the stem growth and organogenesis. Since *AP1* and *CAL* are key genes for FM identity^17,54,55^, we assessed whether *AP1* could also contribute to stem development due to its dominant functions than *CAL*. To test this hypothesis, we examined the RM structure of *ap1-1* mutant during flowering. Unlike the gradual enlarged RM of WT, the RM of *ap1-1* was similar to that of WT at 11 DAG but kept resting state during floral transition and subsequent floral development resulting in tardy stem development, indicating that *AP1* is required for the activation of RM and proper stem development (Fig. 1b; Extended Data Fig.1a). We then used *p35S::AP1-GR;ap1;cal* (*AP1-GR;ap1;cal*) plants, in which a fusion protein of AP1 and the hormone-binding domain of the rat glucocorticoid receptor (GR) was constitutively expressed by the CaMV35S promoter. Treatment with dexamethasone (DEX) activated the AP1-GR fusion protein, thereby enabling synchronized initiation of floral development^20,22^. We noticed that *AP1-GR;ap1;cal* showed markedly shorter stems compared to WT with similar flowering time, indicating that the stem development was impaired in the mutant. Notably, activation of AP1-GR during the vegetative stage was sufficient to rescue the delayed stem initiation of *AP1-GR;ap1;cal* (Fig. 1c,d), indicating that *AP1* played a crucial role in promoting stem initiation. Consistently, the *ap1-1* mutant generated shorter stem than WT (Fig. 1e), probably due to the findings that the RM was inactivated during floral transition, and was significantly shorter after bolting in *ap1-1* than in WT (Fig. 1b, f and g).

Previous studies have shown that stem elongation mainly depends on the growth of the first internode, defined as the region between the shoot apex and the top cauline leaf^6,7,56,57^. We noticed that the first internode of *ap1-1* was shorter than that of WT (Fig. 1e). Time-lapse imaging further revealed that, while the first internode of WT grew rapidly, that of *ap1-1* showed only minimal growth, agreement to the impaired activity of RM in *ap1-1*. Interestingly, the circadian rhythm of inflorescence circumnutation under constant light (LL) conditions clearly observed in WT, was greatly abolished in *ap1-1* (Extended Data Movie 1)^58^. To further investigate this phenotype, we examined rhythmic stem growth under both long day (LD; 12h light/12h dark) and LL conditions. In WT, stem growth rate was faster during the nighttime zeitgeber time (ZT) ZT21-ZT9 compared to daytime (ZT9-ZT21) under both conditions, indicating that the rhythmic stem growth in WT was under strong circadian control. In contrast, this rhythmicity was markedly compromised in *ap1-1* (Fig. 1h,i). Collectively, these findings demonstrated that *AP1* was essential for the activation of stem initiation and its circadian growth after floral transition.

### *TOC1* noncell-autonomously and spatially mediates *AP1* function in stem development

To gain deeper insights into circadian-regulated stem growth, we examined the expression patterns of *CCA1* and *TOC1* in meristematic tissues. *TOC1* and *CCA1* were highly concentrated in early stages FMs and IMs, particularly in the PZs and CZs but less in the RMs. Notably, the expression of *TOC1* was obviously decreased in the IMs and FMs of *ap1-1* compared to WT (Fig. 2a), which was confirmed by RT-qPCR (Extended Data Fig. 1b), showing that *AP1* spatially regulates the expression of *TOC1*. To further investigate their functional roles, we generated CRISPR-Cas9 editing mutants of *CCA1* and *TOC1* in L*er*, designated *cca1-8* and *toc1-17*, respectively (Extended Data Fig. 1c). Both mutants displayed comparable flowering time to that of WT (Extended Data Fig. 1d). Nevertheless, while *cca1-8* showed normal stem development compared to WT (Extended Data Fig. 1e,f), *toc1-17* showed dramatically delayed stem elongation characterized by shorter stems and reduced RMs relative to WT, which was mainly due to the inactive RM of *toc1-17* during floral transition like *ap1-1* (Fig. 1e-g and Extended Data Fig. 1g,h). Time-lapse imaging further revealed delayed apical stem growth and a severely impaired circadian rhythm of inflorescence circumnutation in *toc1-17* (Extended Data Movie 1), indicating that *TOC1* was crucial for both stem initiation and circadian stem growth.

**Fig. 2.**
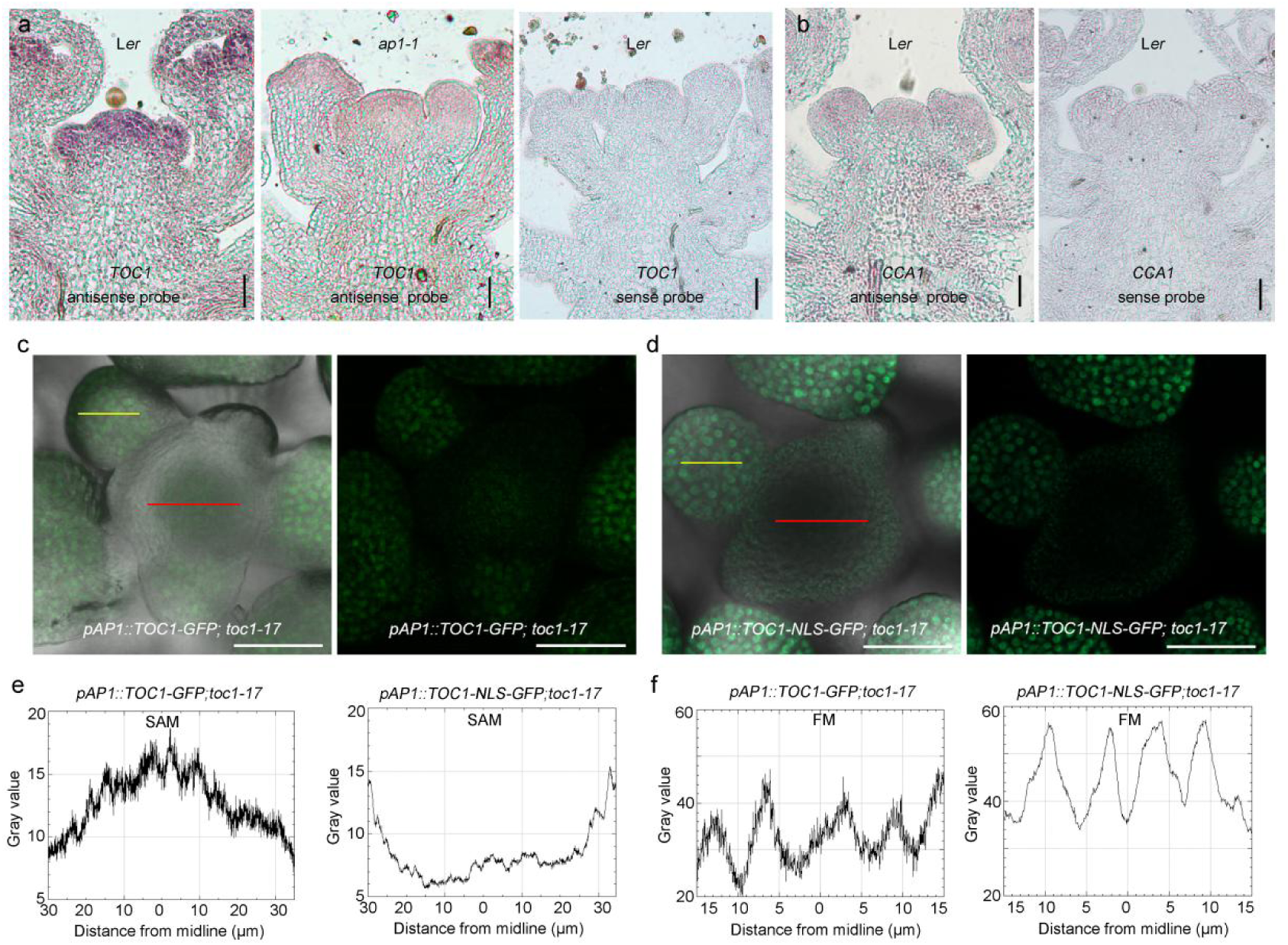
TOC1 noncell-autonomously regulates stem development. (**a** and **b**) *In situ* hybridization to detect expression patterns of *TOC1*(**a**) and *CCA1*(**b**) in the SAMs of indicated plants. Sense probe serves as control. Scale bars: 50 μm. (**c** and **d**) Fluorescence of TOC1-GFP/TOC1-NLS-GFP in IMs and FMs of indicated plants. Fluorescence field alone and merged with bright field were displayed separately. Scale bar: 100 μm. (**e** and **f**) Fluorescence intensity analysis of TOC1-GFP/TOC1-NLS-GFP in the IMs (red line) and FMs (yellow line) of indicated plants in (**c** and **d**).

To assess the genetic relationship between *AP1* and *TOC1* in stem development, we introduced the *toc1-17* into *AP1-GR;ap1;cal* and *ap1-1*, respectively, by crossing, and found that AP1-GR activation failed to restore the stem initiation and elongation but not the floral development in *AP1-GR;ap1;cal;toc1-17* (Fig. 1c,d), suggesting that *TOC1* was required for the roles of *AP1* in stem development but not in floral development. Consistently, the developmental defects of the *ap1-1;toc1-17* were similar to those of *toc1-17* (Fig. 1e-g and Extended Data Movie 1). Strikingly, the expression of TOC1-GFP specifically in FMs by *AP1* promoter (*pAP1::TOC1-GFP;toc1-17*) could fully complement the *toc1-17* phenotype, with detectable GFP fluorescence signal in SAM, indicating a movement of TOC1-GFP from FMs to SAMs (Figs. 1e-i and 2c,d). Conversely, the expression of a nuclear-localized version *TOC1-NLS-GFP* under the same promoter failed to rescue the stem developmental defects of *toc1-17* in *pAP1::TOC1-NLS-GFP;toc1-17* that exhibited no TOC-GFP signals in the SAMs but higher TOC1-NLS-GFP in the FMs than *pAP1::TOC1-GFP;toc1-17* (Fig. 2e,f). These findings indicated that *TOC1* transmited a noncell-autonomous signal from FMs to IMs to spatially regulate stem initiation and circadian growth.

### AP1 interacts with CCA1 and promotes the binding of CCA1 to *TOC1*

To understand the mechanisms underlying AP1-TOC1 module regulated stem development, we first determined whether *AP1* is itself a circadian-regulated gene. We treated the IMs of WT as shown in Fig. 3a as previously discussed^59^, followed by sampling at three biological replicates to examine the circadian expression patterns of the genes of interest. *AP1* was expressed non-rhythmically (Fig. 3b), indicating that it was not a circadian-regulated gene. We then hypothesized that key circadian clock genes such as *CCA1* may mediate the regulation of *AP1* on *TOC1* in this regulatory process. Co-immunoprecipitation assays showed that AP1 physically interacts with CCA1, which was further confirmed by both yeast two hybrid and Luciferase (LUC) complementation assays^60^ (Fig. 3c-e).

**Fig. 3.**
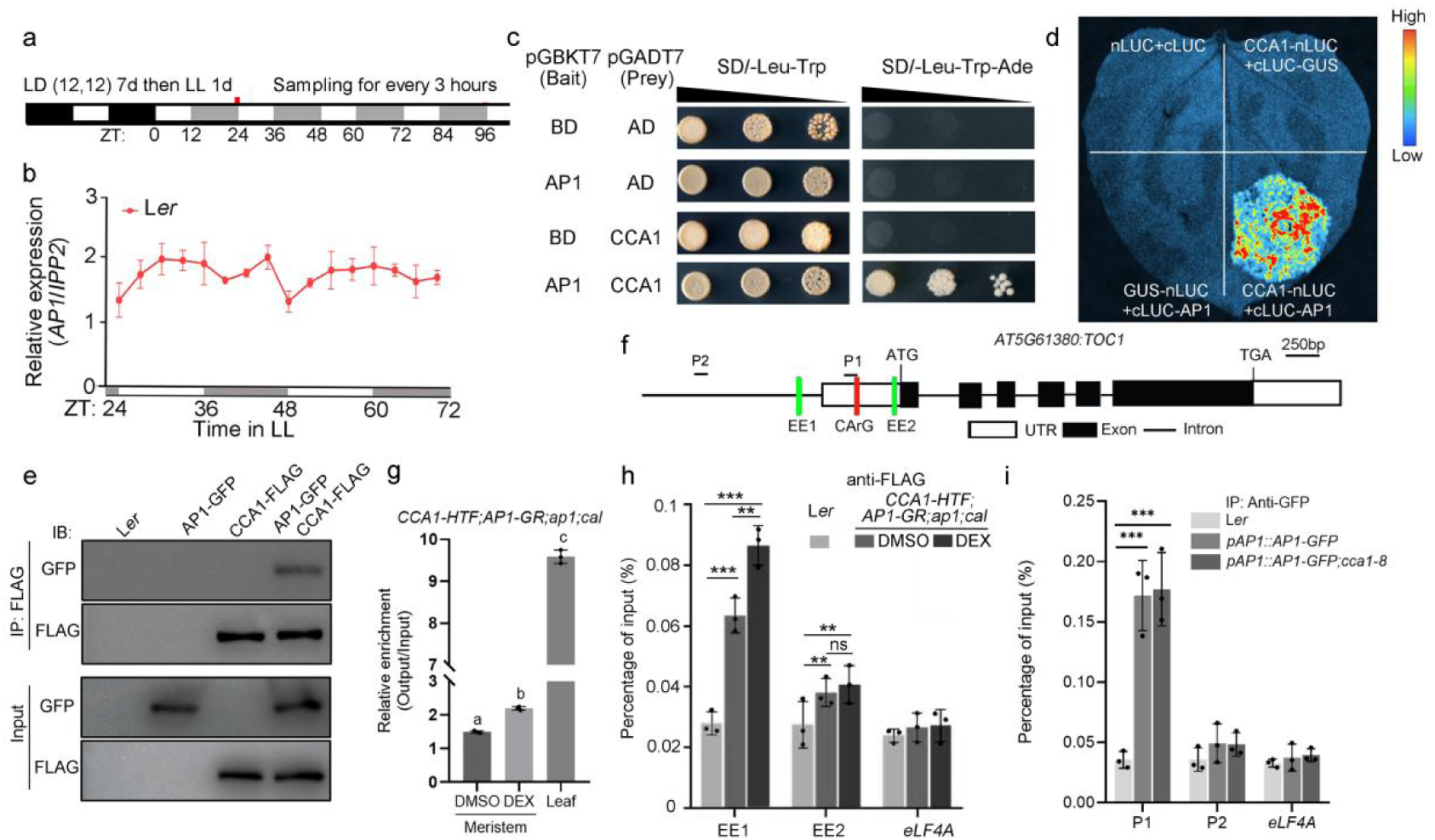
CCA1 interacts with AP1 and regulates *TOC1* in an AP1-dependent manner. (**a**) Schematic diagram of sampling procedure for gene rhythmic expression examination. The plants (indicated in a and Fig4. a-f) were acclimated under light/dark (12/12h) cycles first for 7 days, followed by continuous light exposure. The initial time point was designated as ZT0. (**b**) RT-qPCR of *AP1* expression under the indicated treatments. Data represent mean ± SD of three biological replicates. *IPP2* was used as a normalization control. White and gray bars represent subjective day and night, respectively. (**c**) Yeast two-hybrid system to detect the interaction between AP1 and CCA1. BD, DNA-binding domain; AD, activation domain. (**d**) Split-luciferase assay showing the interaction between AP1 and CCA1 in tobacco leaves. (**e**) Examination of AP1-CCA1 interaction *in vivo* by Co-IP assay. CCA1-FLAG was purified by anti-FLAG antibody from indicated plants and detected AP1-GFP with anti-GFP antibody. IP, immunoprecipitated; IB, immunoblotting. (**f**) Diagram of the *TOC1* locus. White and black rectangles and black lines represent untranslated regions, coding regions and introns, respectively. Three motifs including two EE motifs (green) and one CArG motif (red) and the regions for AP1 binding examination were indicated. (**g**) ChIP-qPCR to examine the binding of CCA1 to *TOC1* in meristems and leaves of indicated plant. ChIP signal was quantified as the percentage of total input DNA by qPCR. Data represent mean ± SD of three biological replicates. Different letters indicate statistically significant differences by one-way ANOVA (Tukey’s multiple comparisons test, P < 0.05). **(h** and **i**) ChIP-qPCR to examine the binding of CCA1 (**h**) and AP1 (**i**) to *TOC1* in the meristems of indicated plants. ChIP signal was quantified as the percentage of total input DNA by qPCR, *eLF4A* served as negative control. The examined loci were indicated in (**e**). Data represent mean ± SD of three biological replicates. Student’s *t*-test (two-tailed); ns denotes not significant; **p < 0.01; ***p < 0.001.

Previous studies revealed that CCA1 specifically bound to the canonical evening element (EE) (AAATATCT) at the transcription start site (TSS) of *TOC1* in seedlings^44^, and AP1, a MADS domain transcriptional factor, binds to CArG-box (CC(A/T)_6_GG) motif ^23,61^. Consistently, two EEs and a CArG-box like motif were identified near the TSS of *TOC1* (Fig. 3f). We were then promoted to examine the binding of CCA1 to *TOC1* by chromatin immunoprecipitation followed by qPCR (ChIP-qPCR) in the seedlings and the IMs of *pCCA1::CCA1-HTF;AP1-GR;ap1;cal* with DMSO (control) and DEX treatment, respectively. Tissues were sampled at ZT48 when CCA1 was highly expressed as shown in Fig.3a. Reproducible enrichment signals and detectable binding occupancy of CCA1 at *TOC1* in the meristematic tissues were observed in DMSO-treated samples, which were further increased in DEX-treated samples (Fig. 3g). Surprisingly, the binding signal of CCA1 to the *TOC1* was significantly lower in the meristematic tissues compared to the seedlings (Fig. 3g)^44^, indicating that CCA1 regulated *TOC1* expression through different mechanisms in meristems and seedlings. Further ChIP-qPCR analysis confirmed that CCA1 bound to both EEs, and that AP1 promoted CCA1 binding specifically to EE1 but not to EE2 (Fig. 3h). Meanwhile, we examined the AP1 binding to TOC1 using the inflorescences of *pAP1::AP1-GFP* and *pAP1::AP1-GFP;cca1-8* by ChIP-qPCR. AP1 was found to bind to the CArG-box like motif, which was independently of CCA1 (Fig. 3i). To confirm the roles of EE and CArG-box in CCA1 and AP1 binding to *TOC1*, we performed yeast one-hybrid assays using a series of site-directed mutations at the EEs and CArG-box within the promoter of *TOC1* as shown in Extended Data Fig. 2a. Both EEs were essential for CCA1 binding to *TOC1*, whereas the CArG-box like motif was required for AP1 binding to *TOC1* (Extended Data Fig. 2b). Collectively, these results suggest that AP1 modulates the regulatory function of CCA1 on *TOC1* expression.

### AP1-CCA1 complex maintains the circadian rhythmicity and function of *TOC1*

To investigate how the AP1-CCA1 interaction regulates *TOC1* circadian expression, we examined the circadian rhythmicity of *CCA1* and *TOC1* in both seedlings and meristems. WT and *AP1-GR;ap1;cal* plants were treated with either DMSO or DEX, as shown in Fig.3a, and leaf and meristem samples were collected every 3 hours starting from ZT24 (Fig. 3a) for RT-qPCR analysis. The circadian expression patterns of *CCA1* and *TOC1* in inflorescences were similar to the respective patterns observed in seedlings (Fig. 4a, b). Under LL conditions, the phase peak of *CCA1* expression in *AP1-GR;ap1;cal* meristems was delayed by approximately 3 hours relative to WT, while the period length and amplitude remained unchanged (Fig. 4c). In the WT shoot apex, *TOC1* was previously found sustaining circadian rhythmicity for over 7 days under LL condition^40^. In contrast, in the *AP1-GR*;*ap1;cal* meristems, the amplitude of *TOC1* expression was reduced at all tested time points, consistent with the *in situ* hybridization result (Fig. 2a), and the circadian rhythmicity progressively diminished, becoming undetectable after 72 hours of free running under LL conditions (Fig. 4d). A similar loss of rhythmicity was also observed in *ap1-1* (Extended Data Fig. 3a), demonstrating that *AP1* was essential for the maintenance of *TOC1* circadian rhythmicity in meristem. Consistently, the activation of AP1 at ZT48 restored the circadian rhythmicity of *TOC1* after one circadian cycle (24 hours), in which the *TOC1* expression was suppressed around ZT72 and elevated again by ZT84 (Fig. 4e; indicated by red and black arrows, respectively). Given that *CCA1* typically reaches the maximum expression in the morning (ZT72) and the minimal expression in the evening (ZT84) ^42^, these results indicated that AP1 interacted with the circadian system by modulating CCA1-mediated regulation of *TOC1*. Specifically, at ZT72, when *CCA1* levels were high, AP1 may recruit CCA1 to the *TOC1* promoter, leading to transcriptional repression. Conversely, at ZT84, when CCA1 levels were extremely low, AP1 may promote *TOC1* expression independently of CCA1 (Fig. 4f), consistent with the known repression role of CCA1 on *TOC1* expression and the transcriptional activation function of *AP1*^24,25,62^. To rule out the possibility that post-transcriptional modifications affect the circadian regulation of *TOC1*, we crossed the *pTOC1::LUC* reporter construct^40^ into *AP1-GR*;*ap1;cal* and monitored *LUC* expression, which reflects *TOC1* transcriptional activity. The *LUC* expression pattern in DEX-treated meristems of *AP1-GR*;*ap1;cal* closely mirrored that of endogenous *TOC1* (Extended Data Fig. 3b), supporting the conclusion that AP1 regulated *TOC1* circadian rhythmicity at the transcriptional level by coordinating with circadian CCA1 accumulation in meristems. Consistently, in *AP1-GR*;*ap1;cal;cca1-8* meristems, activation of AP1 failed to repress *TOC1* expression at ZT72 but still promoted its expression at ZT84 (Fig. 4f), further confirming the role of CCA1 in mediating the time-of-day specific effects of AP1 on *TOC1* transcription.

**Fig. 4.**
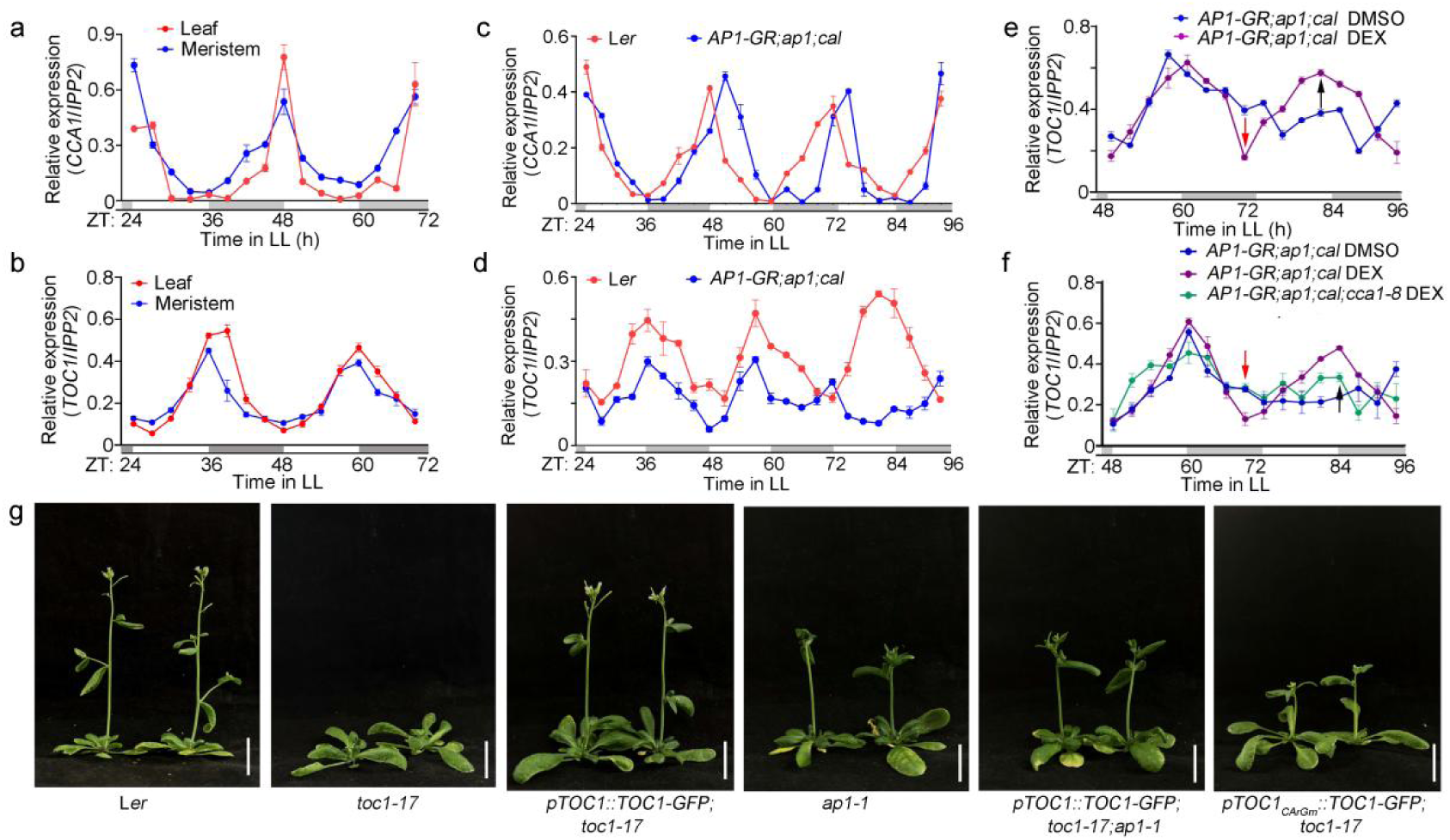
AP1-CCA1 complex maintains circadian expression and functions of *TOC1*. (**a-f**) RT-qPCR to examine the circadian expression patterns of *CCA1* (**a** and **c**) and *TOC1* (**b** and **d**-**f**) in leaves (**a** and **b**) and meristems (**c**-**f**) of indicated plants with or without chemical treatment. Plants were treated with indicated chemicals and the leaves or meristems were sampled at 3-hour interval as shown in Fig. 3a. Data represent mean ± SD of three biological replicates. *IPP2* was used as a normalization control. White and gray bars represent subjective day and night, respectively. Red and black arrows in (**e**) and (**f**) indicated the maximum and minimum expression point of *CCA1*, respectively. (**g**) Morphological observation of indicated plants. Scale bars in (**g**): 2 cm.

To assess the functional significance of AP1 regulated circadian rhythmicity of *TOC1* in stem development, we generated the *pTOC1::TOC1-GFP;toc1-17* construct that could rescue the stem developmental defects of *toc1-17* mutants (Fig. 4g). We then introduced the *pTOC1::TOC1-GFP* into *ap1-1* background. Nevertheless, the expression of *TOC1-GFP* failed to rescue the defects of stem growth and RM size observed in *ap1-1* plants (Fig. 4g and Extended Data Fig. 3c,d), indicating that the functions of *TOC1* on stem development were dependent on *AP1*. Consistently, expression of *TOC1* driven by a site mutated CArG-box in the *TOC1* promoter also failed to complement the stem developmental defects of *toc1-17* (Fig. 4g and Extended Data Fig. 3c,d). These findings suggested that *TOC1* functions downstream of *AP1* to spatially mediate its role in regulating stem development and AP1-CCA1 complex dynamically maintained the circadian rhythmicity and function of *TOC1*.

### AP1 and TOC1 controls stem development through cell development and growth

To further understand the molecular mechanisms underlying *AP1*- and *TOC1*-controlled stem development, we performed transcriptome analysis using inflorescences containing early stage FMs (stages 1-4^63^) and apical stem tissue (sampled at ZT36 when TOC1 was highly expressed as shown in Fig. 3a) from WT, *ap1-1*, *toc1-17* and *pAP1::TOC1-GFP;toc1-17* with three biological replicates (Extended Data Fig. 4a). RNA-seq analysis identified hundreds of different expression genes (DEGs) in *ap1-1* vs. WT, *toc1-17* vs. WT and *pAP1::TOC1-GFP;toc1-17* vs. *toc1-17*, respectively (False Discovery Rate [FDR] < 0.05; Fold Change [FC]≥2) (Extended Data Fig. 4b; Extended Data Table 1.1). Although partially overlapping DEGs were identified in both upregulated and downregulated gene sets of *ap1-1* and *toc1-17* relative to the WT (260/1142 and 147/1140 in *ap1-1*; 260/464 and 147/515 in *toc1-17*), only a small number of DEGs (63 and 69) exhibited opposite expression patterns (Fig. 5a; Extended Data Table 1.1), indicating that *AP1* and *TOC1* acted in similar but distinct regulatory pathways. Gene Ontology (GO) enrichment analysis revealed that biological processes associated with cell development, cell growth, and responses to diverse stimuli were significantly repressed in *ap1-1*. By contrast, pathways involved in reproduction and organismal development were markedly enriched (Extended Data Fig. 4c; Extended Data Table 1.2), indicating that *AP1* played dual regulatory roles in plant developmental processes. Meanwhile, biological processes related to cell growth, as well as specific cell wall modification pathways including pectin catabolic/metabolic processes and galacturonan metabolic process, were downregulated in *toc1-17*, while pathways associated with responses to various stimuli were enhanced (Extended Data Fig. 4d; Extended Data Table 1.3). Consistently, biological processes related to cell development, growth and differentiation, as well as cell wall modification, were simultaneously altered in both *ap1-1* and *toc1-17* (Fig. 5b), in agreement with the stem first internode growth defects observed in the two mutants (Fig.1e; Extended Data Movie 1).

**Fig. 5.**
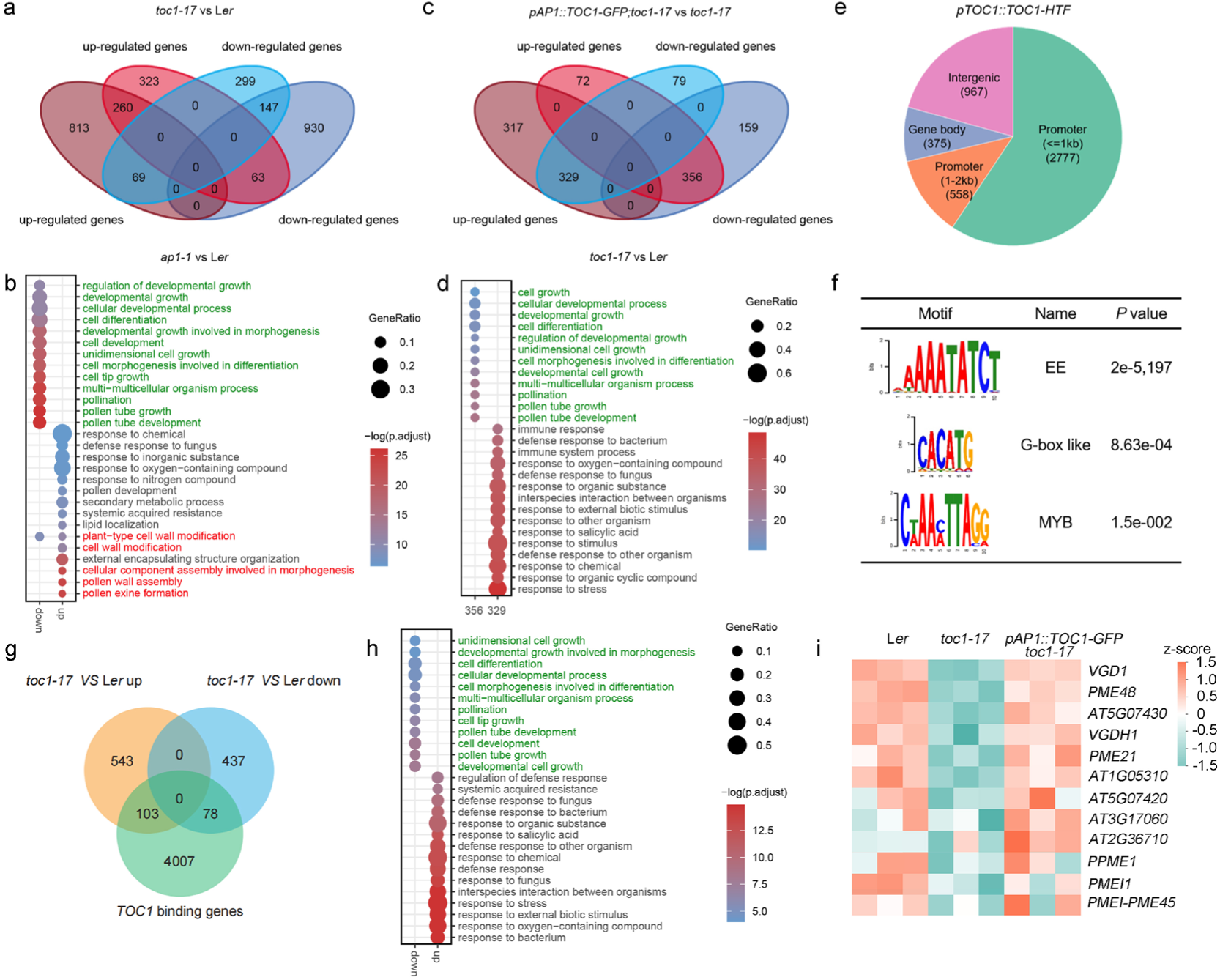
Transcriptomic and ChIP analyses related to Fig. 1. (**a**) Venn diagram showing DEGs between *ap1-1*, *toc1-17*, and L*er* meristem. (**b**) Dot plot showing GO enrichment results of overlapping upregulated genes and downregulated genes in *ap1-1* vs. L*er* and toc1-17 vs. L*er*. (**c**) Venn diagram showing the number of overlapping DEGs between the indicated comparison pair. (**d**) GO analysis of overlapping DEGs in *toc1-17* vs. L*er* down/up-regulated and *pAP1::TOC1* vs. *toc1-17* up/down-regulated genes. (**e**) A genome-wide analysis of the distribution of binding peaks from ChIP-seq data and categorized them into four categories: regions within 1 kb upstream of the transcription start site (Promoter ≤1 kb), regions 1-2 kb upstream of the transcription start site (Promoter 1-2 kb), Gene body, and Intergenic regions. In the *pTOC1::TOC1-HTF* sample, the binding peaks were mainly enriched in the promoter region within 2 kb upstream of the transcription start site (TSS), with the highest enrichment observed in the immediate vicinity of the TSS, accounting for approximately 70% of the total peaks. (**f**) TOC1-binding motifs identified by ChIP-seq analysis. (**g**) TOC1 target genes by combined analysis of ChIP-seq and RNA-seq. (**h**) GO analysis of overlapping DEGs between the indicated comparison pair (**g**). (**i**) Heatmap showing the expression of genes in pectin catabolic and metabolic processes in the indicated plants.

Furthermore, we identified 428 up-regulated and 408 down-regulated genes in *pAP1::TOC1-GFP;toc1-17* compared to *toc1-17*, respectively (Extended Data Fig. 4b; Extended Data Table 1.1). Notably, there was no overlap between up-regulated genes and down-regulated cluster genes within *toc1-17* vs. WT and *pAP1::TOC1-GFP;toc1-17* vs. *toc1-17*. In contrast, a substantial proportion of DEGs showed inverse expression patterns between the two comparisons: 69.1% of the down-regulated genes in *toc1-17* (356 out of 515) were upregulated in *pAP1::TOC1-GFP;toc1-17* (356 out of 428; 83.1%). Similarly, 50.9% of the up-regulated genes in *toc1-17* (329 out of 646) were down-regulated in *pAP1::TOC1-GFP;toc1-17* (329 out of 408; 80.6%) in this comparison (Fig. 5c). GO enrichment analysis of the overlapping DEGs (356 and 329 DEGs) revealed that biological processes associated with cell development, differentiation and growth, as well as pathways involved in responses to diverse stimuli, which were perturbed in *toc1-17*, were substantially rescued in the *pAP1::TOC1-GFP;toc1-17* (Fig. 5d; Extended Data Table 1.4).

Collectively, these results suggested that *AP1*-*TOC1* module noncell-autonomously modulated stem development by regulating gene networks associated with cell development, differentiation and growth, as well as specific cell wall modification processes. These observations align with previous findings showing that gene expression associated with cell wall extension, pectin structural remodeling, and pectin-cellulose interactions predominately control internode elongation in Arabidopsis^56,57^.

### TOC1 occupies distinct genomic loci during reproductive development

To further understand the roles of TOC1 in the reproductive stage, we identified the TOC1 binding sites in the inflorescence of *pTOC1::TOC1-HTF* sampled at ZT36 with three biological replicates by ChIP sequencing (Extended Data Fig. 4e). In total, we identified 4,188 TOC1-binding peaks in inflorescence samples. These peaks were predominantly enriched in the promoter regions proximal to the transcription start site (TSS) of target genes (Fig. 5e and Extended Data Fig. 4e; Extended Data Table 1.5), consistent with previous reports^45,64^. A total of 647 shared TOC1-binding sites were identified in both seedling and inflorescence samples. In contrast, 3,541 and 4,214 sites were uniquely bound by TOC1 specifically in inflorescences and seedlings, respectively (Extended Data Fig. 4f; Extended Data Table 1.5). GO enrichment analysis showed that terms associated with responses to diverse stimuli were enriched among the common TOC1-binding genes shared by inflorescences and seedlings. Inflorescence-specific TOC1 target genes were markedly enriched in organ development and transcription-related processes, whereas seedling-specific targets were preferentially enriched in stimulus response and metabolic pathways (Extended Data Fig. 4g, h; Extended Data Table 1.6). These results indicated that TOC1 exerted distinct regulatory functions in the two tissues.

Previous studies showed that TOC1 acts as one TF binding to the EE (also known as T1ME: TOC1 morning element), G-box (CACTGT) and TBS (TCP binding site: GGCCCA)^45,46,64^. Motif analysis of TOC1-bound sequences identified significantly enriched canonical EE that was bound by both TOC1 and CCA1 (Fig. 5f)^44,45^. Interestingly, a G-box variant (CACATG) and a previously uncharacterized TOC1-binding motif (CWAAMTTAGG) containing a TBS variant (TTAGG) were highly enriched within TOC1-binding peaks (Fig. 5f). These results indicated that TOC1 possesses distinct binding properties and may interact with different co-factors in reproductive tissues. We further deciphered TOC1 target genes by combined analysis of ChIP-seq and RNA-seq, and identifying 103 upregulated and 78 downregulated genes regulated by TOC1 (Fig. 5g; Extended Data Table 1.7). GO enrichment analysis showed that biological processes involved in cell differentiation, development and growth were significantly enriched among TOC1-activated genes. In contrast, terms related to responses to diverse stimuli were markedly enriched in TOC1-repressed genes (Fig.5h; Extended Data Table 1.8), indicating that TOC1 played dual roles in modulating organ development and environmental stimulus responses.

### *VGD1*, a member of the PME family, mediates *TOC1*-dependent regulation of stem development

Previous studies revealed that pectin de-methylesterification triggered by pectin methylesterases (PMEs) is associated with high cell wall extensibility and consequent stem cell identity, meristem maintenance and organ initiation and outgrowth^32,65,66^. Consistent with these findings, we observed that pectin metabolic/catabolic processes were weakened in *toc1-17* (Extended Data Fig. 4d). Further gene expression analysis showed that a group of *PME*s showed decreased expression in *toc1-17,* but restored the expression in *pAP1::TOC1-GFP;toc1-17* as in WT, including the well-characterized *VANGUARD1* (*VGD1*) (Fig. 5i; Extended Data Table 1.9).^32,34^ Protein-Protein Interaction (PPI) network analysis revealed that VGD1 acts as one most important hub protein among the networks regulating stem development (Extended Data Fig. 4i). We further detected similar expression level of *VGD1* in all tested plants at ZT9; however, *VGD1* expression was enhanced in WT at ZT21, which was decreased in the *ap1-1*, *toc1-17* and *ap1-1;toc1-17*, but restored in *pAP1::TOC1-GFP;toc1-17* instead of *pAP1::TOC1-NLS-GFP;toc1-17* (Fig. 6a), consistent with the fast cell growth at nighttime. ChIP-Seq analysis of the *VGD1* locus showed that TOC1-binding to both promoter and 3’UTR regions, in which typical EEs were detected (Fig. 6b). ChIP-qPCR using specific primers targeting TOC1-binding peaks was performed to verify the direct binding of TOC1 to the *VGD1* locus (Fig. 6c, d). Furthermore, we sought to examine its regulatory effect on*VGD1* using a *LUC* reporter assay. The *pVGD1::LUC::3’UTR* construct was co-transformed with *p35S::TOC1* into *Nicotiana benthamiana* leaves. Interestingly, TOC1 significantly enhanced LUC activity (Fig. 6e, f), indicating that TOC1 directly activated *VGD1* expression. To further determine whether the EEs bound by TOC1 mediate TOC1-dependent activation, we generated site specific mutations into these EEs within the LUC reporter system. Mutations of these EEs in both the promoter and 3’UTR regions greatly attenuated the ability of TOC1 to activate *VGD1* expression (Fig. 6e, f). These findings demonstrated that TOC1 directly activated *VGD1* expression, thereby contributing to the maintenance of the expression rhythm of *VGD1*.

**Fig. 6.**
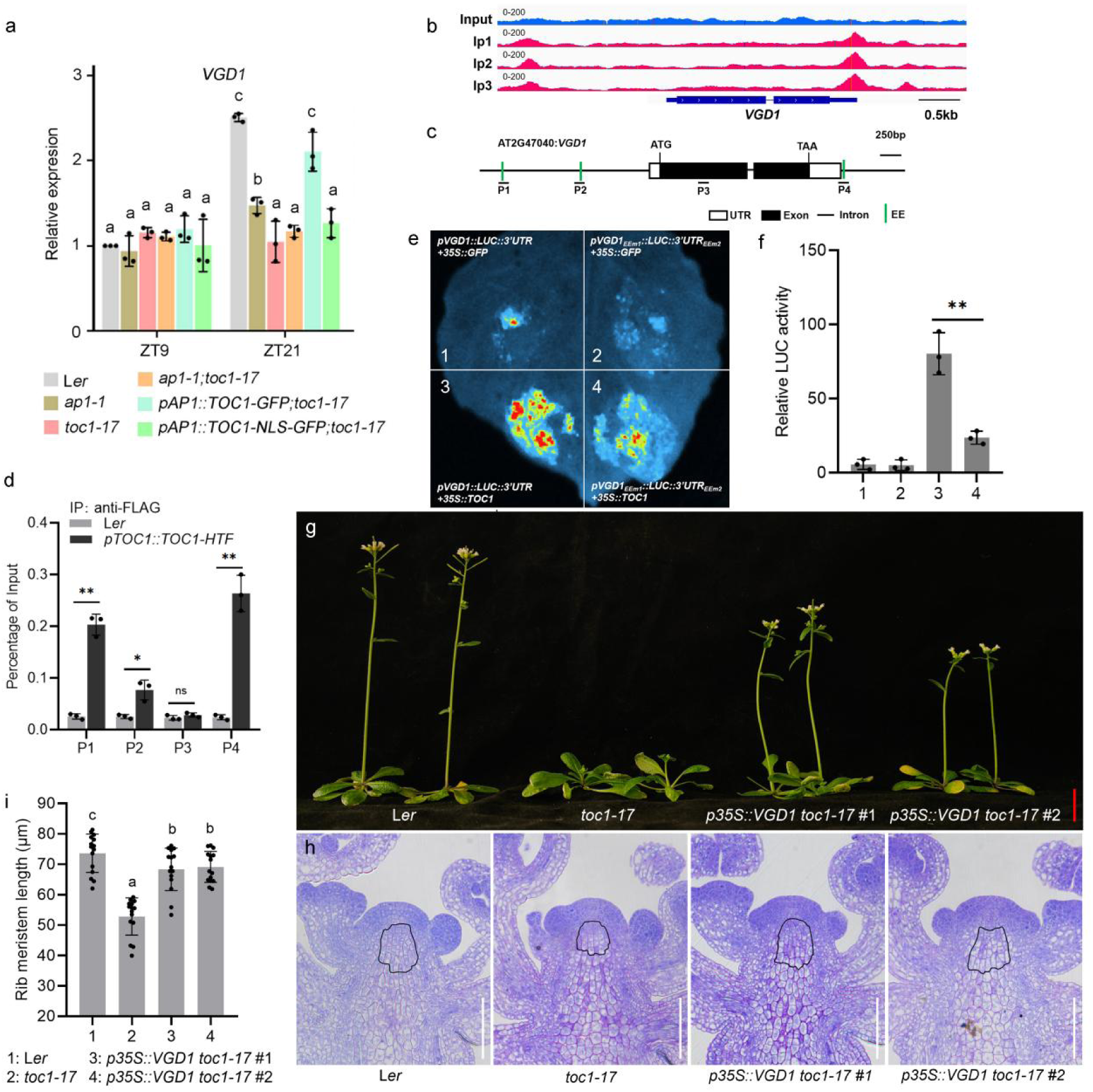
TOC1 directly binds to and regulates *VGD1* to modulate stem development. (**a**) RT-qPCR to examine the expression of *VGD1* in indicated plants at ZT9 and ZT21. Data represent mean ± SD of three biological replicates. *IPP2* was used as a normalization control. The *VGD1* expression level in L*er* at ZT9 was set as 1. (**b**) ChIP-seq analysis revealed a significant binding peak of TOC1 in the upstream region of the *VGD1* gene promoter (The top panel shows the input, and the bottom panel shows three biological replicates of the IP). (**c**) Diagram of the *VGD1* locus. White and black rectangles and black lines represent untranslated regions, coding regions and introns, respectively. (**d**) ChIP-qPCR to examine the binding of *TOC1* to *VGD1* in the meristems of indicated plants. ChIP signal was quantified as the percentage of total input DNA by qPCR. The examined loci were indicated in (**c**). Three EE motifs (green) and the regions for TOC1 binding examination were indicated. Data represent mean ± SD of three biological replicates. Student’s *t*-test (two-tailed); ns denotes not significant; **p < 0.01; ***p < 0.001. (**e** and **f**) LCI system (left) and quantitative analysis of LUC activity (**e**) to examine the direct activation of TOC1 on *VGD1* expression. The *pVGD1::LUC-3’UTR* (1) or *pVGD1_EEm1_::LUC-3’UTR_EEm2_* (2) was transformed with *p35S::GFP* into tobacco leaves, while the *pVGD1::LUC-3’UTR* (3) or *pVGD1_EEm1_::LUC-3’UTR_EEm2_* (4) was transformed with *p35S::TOC1* into tobacco leaves. For quantitative analysis, the LUC activity of *pVGD1::LUC-3’UTR* transformed with *p35S::GFP* was set as 1. Three independent experiments were conducted with similar results. (**g**) Morphological observation of indicated plants. Scale bar: 2 cm. (**h**) Longitudinal sections of shoot apex of indicated plants during reproductive stage showing the length of Rib Meristem (RM). (**i**) Statistical analysis of RM length of indicated plants. Data represent mean ± SD (n=15). Different letters indicate statistically significant differences by one-way ANOVA (Tukey’s multiple comparisons test, P < 0.01).

To investigate the role of *VGD1* in mediating *TOC1*-regulated stem development, we generated *p35S::VGD1; toc1-17* transgenic lines, which exhibited a complementation phenotype of *toc1-17* with respect to stem development (Fig. 6g). Longitudinal sections showed that the delayed activation of the RM in *toc1-17* was rescued by *VGD1* overexpression (Fig. 6h, i). These findings demonstrated that *VGD1* acted as one representative of *PME*s, mediating the role of *TOC1* in regulating circadian stem growth.

### *AP1* homologs have similar function in stem growth in wheat and rice

Plant height, primarily determined by stem development, is a key agronomic trait that significantly influences crop yield.^11,14,67^ To investigate the functions of *AP1* homologs in crops species, we conducted phylogenetic analysis and identified wheat *TaVRN1* and rice *OsMADS14* and *OsMADS18* as homologs of *AP1* (Extended Data Fig. 5a).^68^ *TaVRN1* has been mainly characterized by its role in the vernalization of winter wheat^69–72^, where its expression is induced in apical meristems and leaves to promote flowering^70^. In the absence of vernalization, weak winter wheat cultivar Kenong199 failed to express *TaVRN1* resulting in an inability to complete spike development and initiating stem elongation 4 weeks after germination. However, overexpression of *TaVRN1* in Kenong199 induced both stem elongation and flowering even without vernalization (Fig. 7a). Consistently, following vernalization, *TaVRN1* was activated, leading to stem elongation. Notably, *TaVRN1* overexpressing lines showed longer peduncles and a higher night/day stem elongation ratio compared to Kenong199 (Fig. 7b-e). These results proved that *TaVRN1* was critical for the initiation of stem growth. Common wheat carries three copies of *TaVRN1*, named *TaVRN1-A*, *TaVRN1-B* and *TaVRN1-D*. A dominant (*Vrn*) allele at any one of these loci is sufficient to confer the spring growth habit observed in spring wheat cultivars^73^. Using CRISPR/Cas9 system, we generated loss of function mutants (*Tavrn1-a*, *Tavrn1-b*, *Tavrn1-d* and *Tavrn1-a/d* mutants) in the spring wheat cultivar Fielder (Fig. 7f). All mutants showed reduced plant height, particularly due to the shortened peduncle length, compared to WT, consistent with previous findings (Fig. 7g, h) ^72^. Remarkably, WT showed faster stem elongation during the night compared to the daytime indicating that circadian growth pattern was significantly attenuated in the *Tavrn1* mutants (Fig. 7i). These findings indicated that *TaVRN1* regulated plant height by controlling stem circadian elongation. To evaluate the agronomic potential of *TaVRN1*-regulated plant height, we developed a recombinant inbred line (RIL) population using the spring wheat cultivar Zhongkemai138 and the weak winter wheat cultivar Yumai38 as parents. Without vernalization, Zhongkemai138 displayed earlier heading time and higher plant height compared to Yumai38. After maturation, the peduncle, rather than the other internodes of Zhongkemai138 was significantly longer than that of Yumai38, consistent with differences in parental *TaVRN1* expression levels (Fig. 7j, k). Gene expression analysis and phenotypic observations revealed that RILs with higher *TaVRN1* expression produced longer peduncles and displayed higher night/day stem growth ratio compared to RILs with lower *TaVRN1* expression, resulting in significant variation in plant height across the population under non-vernalized conditions (Fig. 7l-n).

**Fig. 7.**
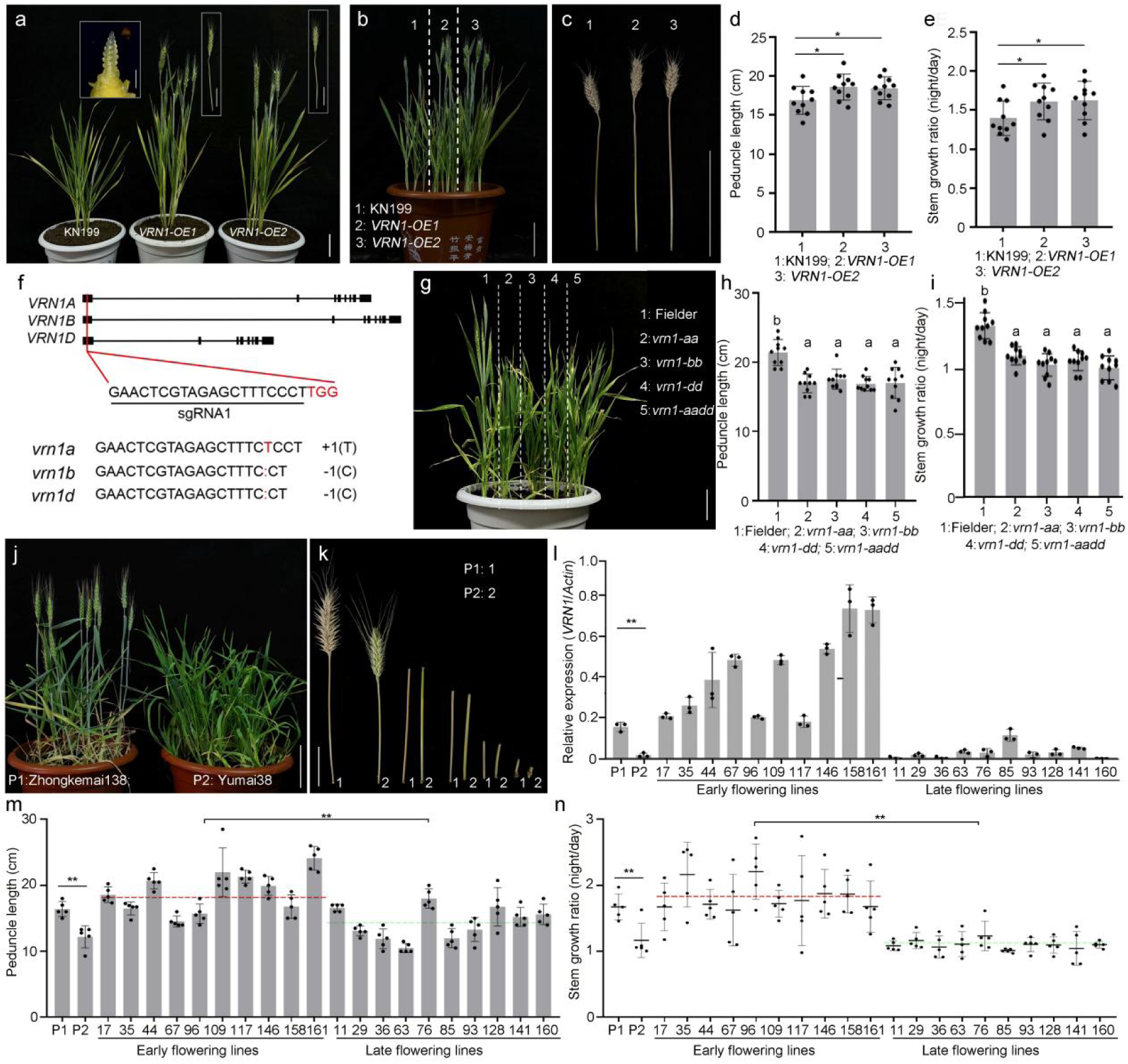
AP1 homologs regulate stem development in wheat. (**a-c**) Morphological observation of KN199, *VRN1-OE1* and *VRN1-OE2* without (**a**) and with (**b** and **c**) vernalization. Scale bars: 5 cm (**a**), 10 cm (**b** and **c**). (**d** and **e**) The peduncle length and stem growth ratio of the KN199, *VRN1-OE1* and *VRN1-OE2* with vernalization. Data represent mean ± SD (n=10). Different letters indicate statistically significant differences by one-way ANOVA (Tukey’s multiple comparisons test, *P* < 0.01). (**f**) Information on the gene editing of different subgenes of *VRN1*. Scale bars: 5 cm. (**g**) Morphological observation of Fielder (WT) and *VRN1* mutants *vrn1-aa*, *vrn1-bb*, *vrn1-dd* and *vrn1-aadd*. Scale bars: 10 cm. (**h** and **i**) The peduncle length (**h**) and stem growth ratio (**i**) of wheat in (**g**). Data represent mean ± SD (n=10). Different letters indicate statistically significant differences by one-way ANOVA (Tukey’s multiple comparisons test, P < 0.01). (**j** and **k**) Morphological observation of wheat Zhongkemai138 and Yumai38 as parents P1 and P2. Scale bars: 10 cm (**j**), 5 cm (**k**). (**l**) RT-qPCR to examine the expression of *VRN1* in the RILs with Zhongkemai138 and Yumai38 as parents. Random early flowering lines and late flowering lines (n=10) were selected with parental lines as control. Data represent mean ± SD of three biological replicates. Student’s *t*-test (two-tailed), **p < 0.01. (**m** and **n**) The peduncle length (**m**) and stem growth ratio (**n**) of indicated wheat accessions in (**i**). Data represent mean ± SD (n = 10). Average of early flowering lines and late flowering lines was marked by red and green lines, respectively. Student’s *t*-test (two-tailed), **p < 0.01.

Previous studies have demonstrated that the rice genes *OsMADS14*, and *OsMADS18* play crucial roles in flowering time and plant architecture^68,74^. To investigate their functions in stem elongation, we generated CRISPR-Cas9-edited mutants of *OsMADS14* and *OsMADS18* in ZH11 (Extended Data Fig. 5b). Under LD conditions in a growth chamber, the plant height of the mutants was significantly reduced compared to that of ZH11. Moreover, while ZH11 displayed stem circadian growth characterized by a higher night/day growth ratio, this rhythmic pattern was disrupted in the mutants (Extended Data Fig. 5c-e). These findings demonstrated that *TaVRN1* and *OsMADS14/18* worked similarly to *AP1* in regulating stem elongation and circadian growth, and suggested that these genes could be candidates for the genetic manipulation of plant height to improve crop yield.

## DISCUSSION

During plant evolution, the reorientation of cell division planes and the establishment of asymmetric cell divisions have promoted the transition from two-dimensional (2D) to 3D growth of plants, ultimately enabling the evolution of the stem development and contributing to the diversification of shoot architecture.^3,75,76^ However, despite the classic morphological and physiological studies of early stem development conducted in the 1950-60s, relative little attention has been given to primary stem growth compared to studies focused on stem secondary growth and vascular development.^2^ In *Arabidopsis thaliana*, the RM remains quiescent during the vegetative stage and is activated to initiate stem development during floral transition. Recent studies revealed that stem growth was stimulated by GA, and locally regulated by *RPL* and its homologs through the activation of boundary-specific target genes.^7,77,78^ Nevertheless, the mechanisms underlying RM activation and stem initiation remain largely unknown. Given that the most distinct morphological change during the floral transition is the generation of FMs rather than leaf primordia, it is reasonable to hypothesize that signals coming from FMs spatially activate RMs and coordinate organogenesis with stem development to establish optimal shoot architecture.

Here, we demonstrated that *AP1* was essential for RM activation, stem initiation, and stem circadian growth during the floral transition (Fig. 1 and Extended Data Fig.1). However, *AP1* expression was excluded from the IMs by *TFL1*^79,80^. Therefore, an intermediary factor was required to spatially mediate the functions of *AP1* in regulating stem initiation and circadian growth. We identified the circadian clock gene *TOC1* as this key mediator. Mutation of *TOC1* disrupted the function of *AP1* on stem initiation and circadian growth, while leaving FM identity and floral development largely unaffected (Fig. 1), suggesting that *AP1-TOC1* module exerted separable roles in stem versus floral development, a conclusion further verified by RNA-seq analysis (Fig. 5a). Consistently, *AP1* maintained both the expression levels and circadian rhythmicity of *TOC1* (Figs. 2 and 4). Notably, expression of TOC1 but not a nucleus-restricted version of TOC1 within FMs was sufficient to rescue the stem developmental defects of *toc1-17*, which was genetically dependent on *AP1* (Fig. 1, 2 and 4). These findings indicated that *TOC1* non-cell-autonomously mediated the functions of *AP1* during stem development. However, the mechanisms by which *AP1* modulates *TOC1* expression in the IM and how TOC1 moves non-cell-autonomously from the FM to the IM remains to be further investigated. Strikingly, homologs of *AP1* in wheat and rice also exhibited similar functions to regulate stem elongation and circadian growth (Fig. 7), consistent with previous findings,^47,68,81,82^ and suggesting potential applications in the regulation of crop height.

Circadian processes are tightly regulated by both external and internal signals with distinct organ- and tissue-specific patterns, which are integrated through the activity of circadian core oscillators^36^. Generally, both CCA1 and TOC1 bind to the EE at target gene loci and directly regulate genes involved in multiple biological processes and stress responses in seedlings. However, genome-wide studies have revealed that TOC1 and CCA1 exert widespread effects on gene expression, extending far beyond the core components of the circadian oscillator^44–46^. This suggests that core oscillator proteins act in a tissue-specific manner via distinct binding motifs or interacting cofactors. For example, EARLY FLOWERING 3 (ELF3), ELF4 and LUX ARRHYTHMO format protein complex controlling hypocotyl growth, while DET1 interacts with CCA1 and LHY mediating transcriptional repression within the plant circadian clock^51,83^. Meanwhile, TOC1 interacts with PHYTOCHROME INTERACTING FACTOR 5 (PIF5), PIF4 and FAR-RED ELONGATED HYPOCOTYL3 to regulate target genes expression^52,64^. In this study, we identified one TOC1-binding G-box variant (CACATG) and a novel TOC1-binding motif (CWAAMTTAGG) in reproductive tissues (Fig. 5f), indicating that TOC1 operates via distinct mechanisms in inflorescences relative to seedlings. Surprisingly, we found that the occupancy of CCA1 at *TOC1* promoter was significantly lower in meristems than in seedlings, but was markedly increased upon AP1 activation, showing that AP1 physically interacted with CCA1 and directed CCA1 binding to *TOC1* promoter (Fig. 3). Considering the circadian accumulation pattern of CCA1, this dynamic represented a sophisticated regulatory mechanism by which AP1-CCA1 coordinately regulated *TOC1* expression: at the accumulation peak of CCA1, AP1 recruited CCA1 to the *TOC1* promoter to repress *TOC1* expression; whereas when CCA1 levels decline, AP1 alone activated *TOC1* expression, thereby maintaining its circadian rhythmicity (Fig. 4e,f). Whether AP1 globally resets circadian-regulated genes expression through coordinated action of CCA1 and TOC1 remains an important question for future investigation.

Previous studies revealed that *TOC1* mediates plant circadian growth^48,50,52,84^. However, the underlying physiological basis remains largely unclear. In this study, we found that *TOC1* was critical for stem initiation and circadian growth by acting downstream of *AP1*. Transcriptome analysis revealed that genes involved in cell wall organization and cell growth showed decreased expression in the *toc1-17* mutant, suggesting that *TOC1* may function as a transcriptional activator (Figs. 1, 2 and 4). Although TOC1 is generally characterized as a transcriptional repressor ^45,46^, our target gene analyses revealed that TOC1 tends to activate genes associated with developmental processes while repressing those involved in diverse stress responses (Fig. 5g, h), indicating that TOC1 functions as both a transcriptional activator and repressor. Recent evidence has reported that OsTOC1 acts as an activator to directly activate a target gene in pathogen resistance in rice^47^. Consistent with this, we demonstrated here that TOC1 directly activated *VGD1* expression to maintain its rhythmic expression (Fig. 6a-d). These findings suggested that TOC1 may act either as repressor or activator, potentially depending on interactions with specific co-regulators. A similar dual functionality has been reported for the transcriptional repressors CCA1 and AP1, which are also capable of activating specific target gene expression^23,85^. Thus, elucidating the context-dependent regulatory roles of TOC1, CCA1 and AP1 in different developmental stages represents an important direction for future research. Recent studies reported that the cell wall properties are critical for stem cell fate within the SAM of Arabidopsis, in which PMEs play important roles in modifying the methylesterification and de-methylesterification of pectins to modulate the chemical and mechanical properties of the cell wall^32,33^. In this study, we revealed that *TOC1* modulated stem development primarily by targeting the expression of *PME*s, suggesting that PMEs-mediated dynamic alterations in pectin properties contribute to circadian-regulated stem growth. Therefore, we delineate an elaborate molecular and genetic pathway through which the AP1/CCA1-TOC1-PMEs cascade spatiotemporally modulates stem development.

## METHODS

### Plant materials

*Arabidopsis thaliana* ecotype Landsberg *erecta* (L*er*) was used as the wild type (WT) reference in this study. The following previously characterized Arabidopsis lines were also used: *ap1-1*^86^, *35S::AP1-GR;ap1;cal*^22^, *pAP1::AP1-GFP*^87^ and *pCCA1::CCA1-HTF*^88^. For wheat experiments, the bread wheat (*Triticum aestivum L.*) cultivars KN199 and Fielder were used as wild type controls in this study, and the transgenic line *VRN1-OE* has been previously described^89^. For rice experiments, the japonica rice (*Oryza sativa L*.) variety Zhonghua11 (ZH11) was used as the wild type reference.

### Plant growth conditions and treatments

All *Arabidopsis thaliana* plants were cultured either in a greenhouse or in a growth chamber (BPC600H, JIUPO, China) maintained at 22°C under a 12 h light/12 h dark (LD) photoperiod or under constant light (LL) conditions. Healthy wheat seeds were surface sterilized using 10% H_2_O_2_ and then germinated on filter paper moistened with distilled water, and incubated in darkness at 25 °C for 1 day. Germinated seeds were then transplanted into pots and grown under a 12 h light/12 h dark cycle at 25℃ in a greenhouse. For genotypes requiring vernalization, seedlings were subjected to cold treatment at 4°C for 3-4 weeks before to be transferred to greenhouse conditions. Rice seeds were directly transplanted into pots and grown under a 12 h light/12 h dark cycle at 28℃ in a greenhouse. *Nicotiana benthamiana* plants, five weeks old and grown in soil under a 12 h light/12 h dark cycle at 22℃ were used for transient expression assays.

### Plasmid construction

The promoter and coding sequences (CDS) of *TOC1*, *CCA1*, *AP1* were amplified using L*er* genomic DNA or cDNA as the template. The resulting fragments were then cloned into the *pEarleyGate301* or *pMDC107* vectors via restriction digestion and ligation to generate the following in-frame fusion constructs: *pTOC1::TOC1-GFP* and *pAP1::TOC1-GFP,* respectively. To generate a mutated version of *pTOC1::TOC1-GFP*, site directed mutagenesis and PCR amplification was performed to obtain *pTOC1_CArGm_::TOC1-GFP*. In addition, the construct *pAP1::TOC1-NLS-GFP* was generated by inserting a nuclear localization signal (NLS) sequence downstream of the *TOC1* coding region in the *pAP1::TOC1-GFP* background. Both constructs were transformed into the *toc1-17* mutant background. In addition, the CDS of *TOC1* was amplified using L*er* cDNA as a template and cloned into *pMDC83* vector to generate the *p35S::TOC1-GFP* construct. To construct *pVGD1::LUC-3’UTR*, a 2 kb sequence upstream of the *VGD1* start codon and a 0.5 kb sequence downstream of the stop codon were amplified and cloned into the *pEarleyGate301* vector, followed by the insertion of the LUC reporter gene downstream of the *VGD1* coding region. Site directed mutagenesis of this construct (*pVGD1::LUC-3’UTR*) was then performed to generate *pVGD1_EEm1_::LUC-3’UTR_EEm2_*. All constructs were introduced into *Agrobacterium tumefaciens* strain *GV3101* for downstream assays. To create the yeast two-hybrid constructs, the *CCA1* and *AP1* coding sequences were amplified using L*er* cDNA as a template and separately cloned into *pGADT7* (activation domain, AD) and *pGBKT7* (binding domain, BD). The plasmid integrity was confirmed by Sanger sequencing. To generate the *CCA1-nLUC* and *AP1-cLUC* constructs, the coding sequences of *CCA1* and *AP1* were amplified using L*er* cDNA as a template and separately cloned into *pENTR/D-TOPO* vector. The destination plasmids were confirmed by Sanger sequencing. The plasmid was linearized and recombined into the destination vectors *pCAMBIA1300-cLUC* and *pCAMBIA1300-nLUC* using the Gateway LR Clonase II enzyme mix (Invitrogen) following the manufactureŕs protocol. All primers used for vector construction are listed in Extended Data Table 1.10.

### Generating the *cca1-8* and *toc1-17* mutants

To generate targeted mutations in *CCA1* and *TOC1*, guide RNAs (gRNAs) were designed to target the second exon of *CCA1*, and the second and third exons of *TOC1*. The gRNA-specific primers (listed in Extended Data Table 1.10) were annealed and ligated into the *pHEE401* vector, following digestion by *Bsa*I^90^. The resulting constructs, CRISPR-CCA1 and CRISPR-TOC1 were independently transformed into L*er* plants via Agrobacterium mediated transformation. CRISPR/Cas9 yielded a 59 bp deletion in *cca1-8* and a 310bp deletion in *toc1-17*, respectively. The deletions resulted in frameshift mutations that are predicted to disrupt the open reading frame and produce nonfunctional truncated proteins.

### *in situ* hybridization

*in situ* hybridization was performed as previously described ^91^. The inflorescences of Arabidopsis were collected and fixed with 4% (w/v) paraformaldehyde (PFA), followed by a graded ethanol series for dehydration and then paraffin embedding. Tissue sections were cut at a thickness of 8μm for *in situ* hybridization analysis. RNA probe templates were amplified from cDNA using gene-specific primers containing T7 or T3 promoter sequences at their 5’ ends. Digoxigenin (DIG)-labeled RNA probes were synthesized *in vitro* using T7/T3 RNA polymerase (Roche, 10881767001) and digoxigenin-UTP (Roche, 11277073910).

### Histological sectioning

For semithin sectioning, the inflorescences of Arabidopsis were fixed using FAA solution (50% ethanol, 5% glacial acetic acid, and 3.7% formaldehyde). Samples were then subjected to vacuum infiltration for 30 minutes to enhance fixation, followed by graded ethanol dehydration (30%, 50%, 70%, 95%, and 100%, 30 minutes each step). Safranin O (0.05%) was added to the 95% and 100% ethanol solutions to stain the tissues. Following dehydration, samples were gradually infiltrated and embedded in resin blocks using a resin embedding kit (Kulzer Technik, Technovit 7100) according to the manufactureŕs instructions. Embedded tissues were sectioned to a thickness of 1µm for microscopic analysis.

For paraffin sectioning, aboveground tissues from 13-day-old *Arabidopsis* seedlings were harvested and fixed in FAA solution (50% ethanol, 5% glacial acetic acid, and 3.7% formaldehyde) under vacuum infiltration for 30 minutes, followed by fixation at room temperature for an additional 24 hours. Samples were then dehydrated through a graded ethanol series (50%, 70%, 95%, and 100% ethanol, 30 minutes per step). Tissue infiltration was performed using a xylene substitute (Histoclear) in increasing concentrations: 30% Histoclear/70% ethanol, 65% Histoclear/35% ethanol, and 100% Histoclear (30 minutes each step). Following complete infiltration, tissues were embedded in paraffin. Embedded samples were sectioned at a thickness of 6 µm using a rotary microtome (Leica RM2235, Leica, Shangai).

### Confocal microscopy

The inflorescences of Arabidopsis was excised, and the mature floral organs along with surrounding primordia adjacent to the SAM were carefully removed using a fine needle under a stereomicroscope. Fluorescence imaging was performed using a Leica Stellaris 5 confocal laser scanning microscope. Imaging was performed sequentially through GFP (excitation 488 nm/emission 507 nm), supplemented with additional bright-field observations when required. Bright-field images were converted to grayscale conversion in LAS X software prior to fusion with green channel signals (GFP). Identical fluorescence intensity settings were maintained across experimental replicates. Imaging parameters included the use of a 20× objective lens (numerical aperture (NA) 0.75), and a Z-step size of 2µm.

### ChIP assay

The chromatin immunoprecipitation (ChIP) assay was performed as previously described ^92^. To investigate the chromatin-binding activity of AP1, CCA1 and TOC1 proteins, SAM tissues were collected from bolted Arabidopsis plants of the following genotypes: L*er*, *pAP1::AP1-GFP*, *pAP1::AP1-GFP;cca1-8*, *pCCA1::CCA1-HTF;AP1-GR;ap1;cal*, and *pTOC1::TOC1-HTF;AP1-GR;ap1;cal*. Experimental and control groups were established according to the study design. SAM tissues (2g) were ground into a fine powder in liquid nitrogen and homogenized in M1 buffer (10mM phosphate buffer pH 7.0, 0.1M NaCl, 10mM β-mercaptoethanol, 1M hexylene glycol, protease inhibitor cocktail, 1mM PMSF). Cross-linking was performed with 37% formaldehyde for 10min, followed by filtration through Miracloth. The homogenate was centrifuged at 2,000 × g to pellet nuclei. The nuclear pellet was then sequentially washed with M2 buffer (M1 buffer supplemented with 10mM MgCl₂ and 0.5% Triton X-100) and M3 buffer (M1 buffer supplemented without 1M hexylene glycol), and resuspended in nuclear lysis buffer (50mM Tris-HCl pH 8.0, 10mM EDTA, 1% SDS, protease inhibitor cocktail). Chromatin was then sheared to 200–500 bp fragments by sonication. Chromatin complexes were immunoprecipitated using anti-FLAG or anti-GFP antibodies. Precipitated DNA was purified for subsequent ChIP-qPCR or ChIP-seq analyses. ChIP-qPCR primers are listed in Extended Data Table 1.10.

### Co-IP assay

Homozygous plants of *pAP1::AP1-GFP×pCCA1::CCA1-HTF* were grown for 4 weeks. Inflorescences were harvested and immediately ground into a fine powder in liquid nitrogen. The powdered tissue was resuspended in extraction buffer (50mM Tris-HCl pH 7.5, 150mM NaCl, 0.1% NP-40, 10% glycerol, 1mM PMSF, and 1×protease inhibitor cocktail) and incubated for 40 min with gentle rotation at 4°C. Following incubation, the mixture was centrifuged twice at 21,000 × g for 10min at 4°C. The resulting supernatant was precleaned by incubation with 30μL Anti-FLAG (M2 Affinity gel, Sigma A2220) for 6 hours at 4°C, and washed at least five times with extraction buffer. The protein complexes were then mixed with 2×SDS loading buffer and analyzed by immunoblotting with anti-FLAG and anti-GFP antibodies, respectively ^92^.

### Yeast two-hybrid assay and Yeast one-hybrid assay

The yeast two-hybrid assay was performed using the Matchmaker GAL4 Two-Hybrid System 3 (Clontech) according to the manufacturer’s instructions as previously described ^92^. The indicated constructs, along with the appropriate empty vector controls were transformed into yeast strain Y2HGold using the polyethylene glycol/lithium acetate (PEG/LiAc) method. Transformants were cultured on SD medium without Leu and Trp (SD/–Leu, –Trp) for selection and subsequently transferred to SD/–Ade,–His,–Leu,–Trp for 3 days to assess protein-protein interactions. Plates were incubated at 30°C in the dark for 3 days (on SD/–Leu,–Trp) and 7 days (on SD/–Ade,–His,–Leu,–Trp).

For the yeast one-hybrid assay, the indicated constructs, and the corresponding empty vectors were transformed into yeast strain Y1HGold using the PEG/LiAc method. Transformed yeast cells were plated on the SD/-Trp,-Leu deficient medium and incubated at 30℃. For induction assays, yeast cultures were grown on SD medium lacking Leu, with or without the addition of ABA for 3 days at 30°C.

### Plant height measurement

After one week of acclimation, mutant plants were grown under controlled environmental conditions with a 12h light/12h dark photoperiod (light from 9:00 to21:00, dark from 21:00 to 9:00). A long bamboo stick was inserted adjacent to each plant to serve as a reference for stem height determinations. The stem height was marked on the bamboo stick at 9:00 and 21:00 each day. Growth increments during the day and night periods were then calculated to compare stem elongation among different mutant genotypes.

### LCI assay

The constructed nLUC and cLUC plasmids were transformed into *A. tumefaciens* strain GV3101. *Nicotiana benthamiana* leaves were transiently transformed by infiltrating the Agrobacterium suspension into the abaxial side of the leaves. Following infiltration, the plants were kept in the dark for 10 hours. After 48 hours, the leaves were irradiated with firefly luciferase substrate buffer, and luminescence signals were captured using an imaging system to assess luciferase fluorescence intensity as previously described^93^.

### Luciferase reporter system assay

The constructs *pVGD1::LUC::3’UTR*, *pVGD1_EEm1_::LUC::3’UTR_EEm2_*,and *p35S::TOC1-GFP* were transformed into *A. tumefaciens* strain GV3101. Reporter constructs were co-infiltrated with transcription factors constructs into *N. benthamiana* leaves by syringe infiltration on the abaxial side. After infiltration, plants were incubated in the dark for 10 hours. After 48 hours, the leaves were irradiated with firefly luciferase substrate buffer, and luminescence signals were captured using an imaging system to assess. luciferase fluorescence intensity as previously described^93^.

### Time-lapse photography and video processing

Time-lapse imaging was conducted at 20-minute intervals over a 48-hour period using a Nikon D750 DSLR camera (Nikon Corporation, Tokyo, Japan) mounted within a climate-controlled growth chamber maintained at 22 ± 0.3°C under continuous full-spectrum illumination (150 μmol·m⁻²·s⁻¹). Post-acquisition video processing was executed through Adobe Creative Cloud 2017 software. Image sequences were stabilized and enhanced in After Effects CC 2017 (v14.2) followed by temporal alignment, compression, and export using Premiere Pro CC 2017 (v11.0). Final time-lapse videos were encoded in H.264format at 30 frames per second (fps), with embedded timestamp metadata.

### RNA extraction and quantitative RT-PCR

Inflorescences or SAMs were excised from whole plants for analysis. Total RNA was isolated from the samples with RNAiso Plus (TaKaRa) according to the manufacturer’s instructions. Residual genomic DNA was removed with DNaseI (Roche). For transcriptome sequencing or reverse transcription, complementary DNA (cDNA) was synthesized using the RevertAid First Strand cDNA Synthesis Kit (TaKaRa). Quantitative real time PCR (RT-qPCR) was performed in three biological replicates using SYBR Green based qPCR Master Mix (Vazyme) on a Bio-Rad CFX-96 Real-time PCR Detection System. The gene expression level in each sample was normalized to *IPP2* or *Actin* as internal reference genes. All primers used for RT-qPCR are listed in Extended Data Table 1.10.

### Pre-processing of RNA-seq data and Identification of DEGs

The raw sequencing reads were processed with fastp v0.20.0^94^ to remove low-quality based and adapter sequences, applying default filtering criteria. The quality of the resulting clean reads was assessed using FastQC to ensure high base quality and the absence of adapter contamination. The resulting clean reads were aligned to the *Arabidopsis thaliana* TAIR10 reference genome with HISAT2 v2.2.1^95^. Gene-level read counts were quantified with FeatureCounts v2.0.3^96^. To adjust for variations in sequencing depth across samples, the read count matrix was normalized using the trimmed mean of M-values (TMM) method as implemented in the edgeR package^97^. Genes with log₂ counts per million (log₂CPM) values greater than 1 in at least three samples were considered to be expressed. The resulting normalized count matrix and corresponding log₂CPM values were then utilized for subsequent analyses.

Differential gene expression analyses of the RNA-seq data were performed using the edgeR package. The selected treatment group and control group samples (T vs. C) were compared to identify DEGs. Genes exhibiting a false discovery rate (FDR) below 0.01 and an absolute log₂ fold change (log₂FC) greater than 1 were considered significantly differentially expressed.

### ChIP-seq data analysis

Raw reads were pre-processed using the same pipeline as previously described for time-series transcriptome data preprocessing.

Peaks were called with MACS2 (v2.2.9.1) under stringent criteria: peaks were retained only if they exhibited a width <1,000 bp, fold enrichment >5, and were reproducibly detected in at least two of three biological replicates (q-value <0.01). All identified peaks were annotated using R-package “ChIPseeker” to assign peaks to genomic features (promoter, exon, intron, intergenic) and to visualize peak distributions relative to TSS and TTS. In this analysis, a consensus peak set was created using peaks detected in ≥2 samples. For de novo motif discovery, Peak summit regions (±200 bp) were extracted and analyzed with the MEME Suite (v5.5.2) with default zero or one occurrence per sequence (zoops) parameters. Visualization of chromatin interaction patterns was conducted in IGV (v2.18.4) using merged signal tracks.

### Protein-protein interaction network analysis

A protein-protein interaction (PPI) network analysis on all DEGs identified in *toc1-17* VS. L*er*. Genes with established interactions were retained for network visualizationand annotated *VGD1* as a direct target gene of TOC1 identified by ChIP. This analysis revealed that the interaction network was primarily composed of the PME family and other cell wall-related gene families.

## Accession codes

The ChIP-seq and RNA-seq data have been deposited in the China National GeneBank DataBase (CNGBdb) database under the accession number PRJCA064232.

## Accession Numbers

Sequence data for the genes in this article can be found in the Arabidopsis Genome Initiative or GenBank/EMBL databases under the following accession numbers: AP1: AT1G69120, CAL: AT1G26310, CCA1: AT2G46830, eLF4A: AT3G13920, IPP2: AT3G02780, PME21: AT3G05610, PME48: AT5G07410, PMEI1: AT1G48020, PMEI-PME45: AT4G33230, PPME1: AT1G69940, TOC1: AT5G61380, VGD1: AT2G47040, VGDH1: AT2G47030, OsMADS14: LOC_Os03g54160, OsMADS18: LOC_Os07g41370, TaActin: TraesCS5A02G124300, TaVRN1-A: TraesCS5A02G391700, TaVRN1-B: TraesCS5B02G396600, TaVRN1-D: TraesCS5D02G401500.

## ACKNOWLEDGMENTS

We thank Dr. Robert Sablowski, Wenhao Yan, Xuemei Chen, Yuling Jiao, and Zhong Zhao for the valuable discussion and advice, and Dr. Lei Wang and Shuanghe Cao for sharing *pTOC1::LUC* and *VRN1-OE* lines, respectively. This work was supported by the National Natural Science Foundation of China (grants U24A20391 and 32270340 to X.L., 32300304 to H.Z.), Hebei Natural Science Foundation (grants C2025205068 to X.L., C2024205036 to L.G., C2024205009 to H.Z. and C2025111010 to D.Z.), and the Seed Science and Technology Innovation Team Project of Shijiazhuang (232490472A to X.L.), Hebei Education Department (grant JCZX2025026 to X.L.), startup funds from Hebei Normal University (L2021B19 to X.L.).

## AUTHOR CONTRIBUTIONS

X.L. and H.Z. conceived the project, designed the experiments, and wrote the manuscript. J.X. and H.Z. performed tissue sampling, gene expression analysis such as RT-qPCR and *in situ* hybridization, Confocal analysis and histological section. Y.Y. and C.J. performed the RNA-seq and ChIP-seq analysis, stem circadian growth analysis and time-lapse imaging under the directions of X.X. and Q.X.. J.Q., Y.Z., D.Z., M.J., and L.G. performed genetic and transgenic experiments, phenotypic analyses, protein-protein interaction, yeast one hybrid and transcriptional activity analysis. Y.L., and X.F. conducted functional analysis of *AP1* homologs in wheat and rice. Y.H. participated in the wheat *AP1* homolog analysis. P.G. analyzed and discussed the results and wrote the manuscript with X.L. and H.Z.

---

**Extended Data Fig.1.**
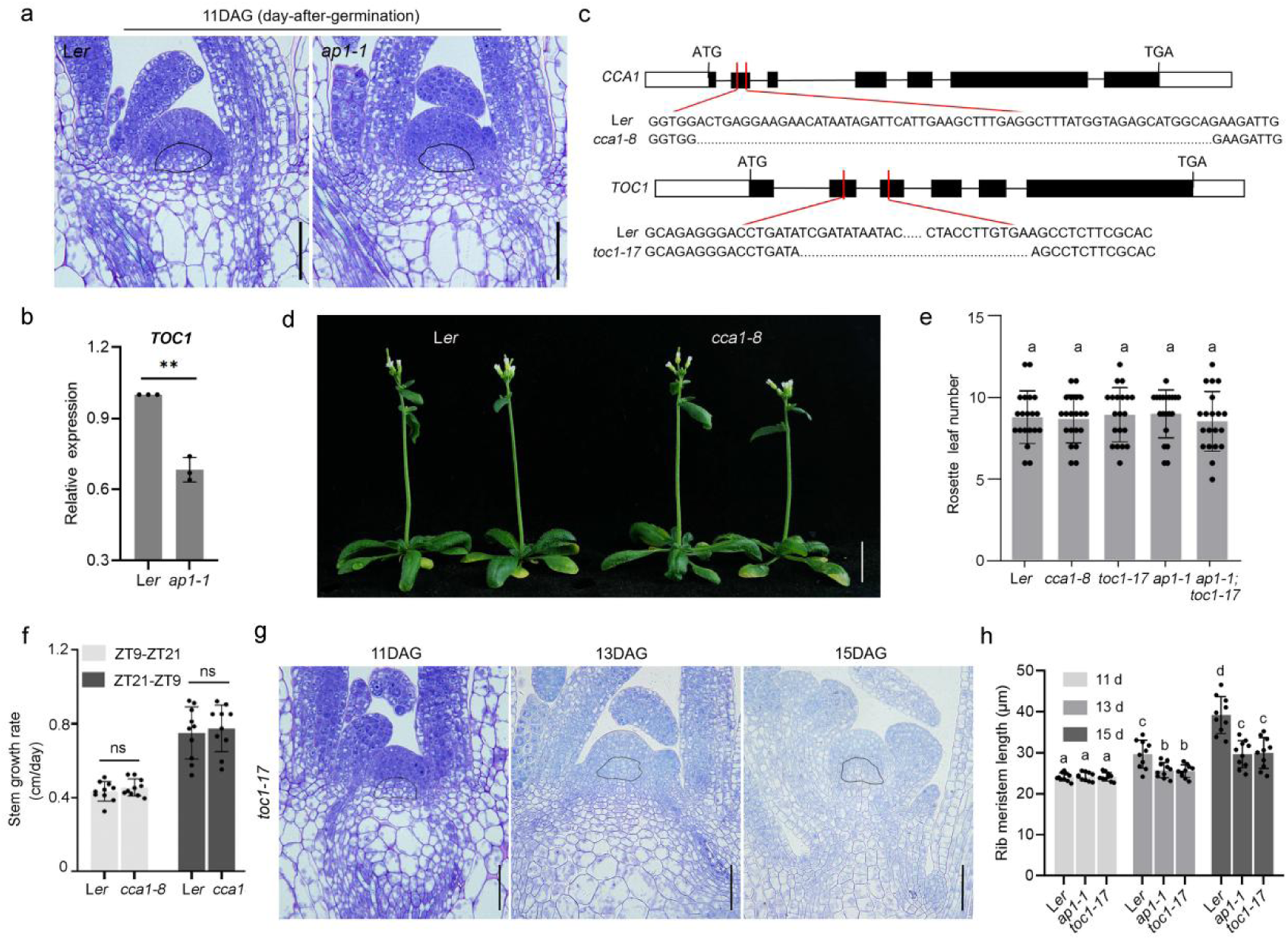
AP1 and TOC1 initiates stem development. (**a**) Longitudinal sections of shoot apex of indicated plants at 11DAG. Scale bars: 50μm. (**b**) RT-qPCR to examine the expression of *TOC1* in the meristems of indicated plants. (**c**) Diagram of the *CCA1* and *TOC1* locus. Gray and black rectangles and black lines represent untranslated regions, exons, and introns, respectively. *cca1-8* and *toc1-17* mutants had one DNA fragment deleted in each (shown in red), 59 bp and 310 bp were deleted respectively. (**d**) Statistical analysis of the rosette leaf number of indicated plants. Data represent mean ± SD. (n=20). Different letters indicate statistically significant differences by one-way ANOVA (Tukey’s multiple comparisons test, P < 0.01). (**e**) Morphological observation of L*er*, *cca1-8* plants. Scale bar: 2 cm. (**f**) The stem growth rate in night and day of L*er* and *cca1-8* under LD condition (12h light/12h dark). ZT, zeitgeber time. Data represent mean ± SD (n=10). Statistical significance was determined by Student’s *t*-test (two-tailed); ns denotes not significant. (**g**) Longitudinal sections of shoot apex of indicated plants. Scale bars: 50 μm. (**h**) Statistical analysis of RM length of indicated plants. Data represent mean ± SD (n=15). Different letters indicate statistically significant differences by one-way ANOVA (Tukey’s multiple comparisons test, P < 0.01).

**Extended Data Fig.2.**
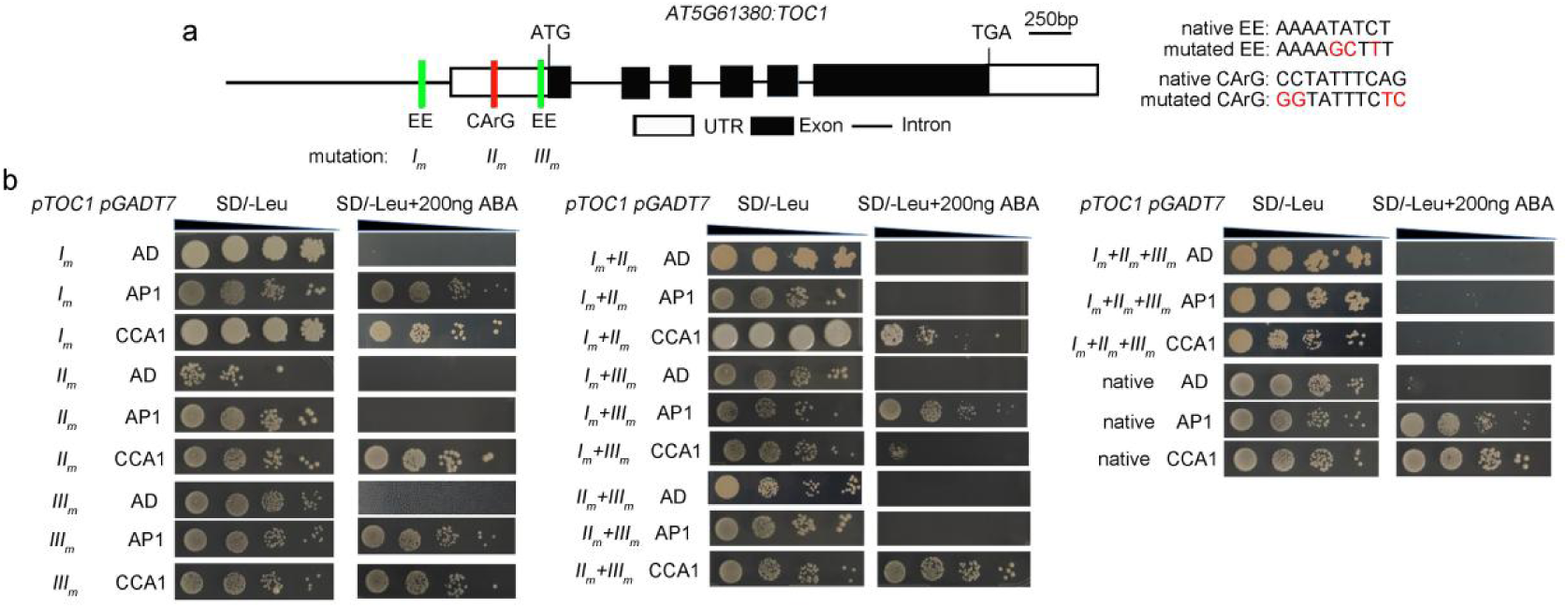
Yeast one-hybrid assays to examine the binding of AP1 and CCA1 to the *TOC1* promoter. (**a**) The diagram of the *TOC1* locus is located above with white and black rectangles and black lines representing untranslated regions, coding regions, and introns, respectively. Three motifs including two EE motifs (green) and one CArG motif (red) were indicated at the promoter of *TOC1*, and the mutated bases were indicated in red on the right. The mutations in each of the three motifs are denoted as *I_m_*, *II_m_*, and *III_m_*, respectively (up panel). (**b**)The target DNA sequences were integrated into the yeast genome with reporter genes and the AP1 or CCA1 proteins were expressed in yeast cells and screened on plates containing SD/-Leu or SD/-Leu+200 ng ABA medium (down panel).

**Extended Data Fig.3.**
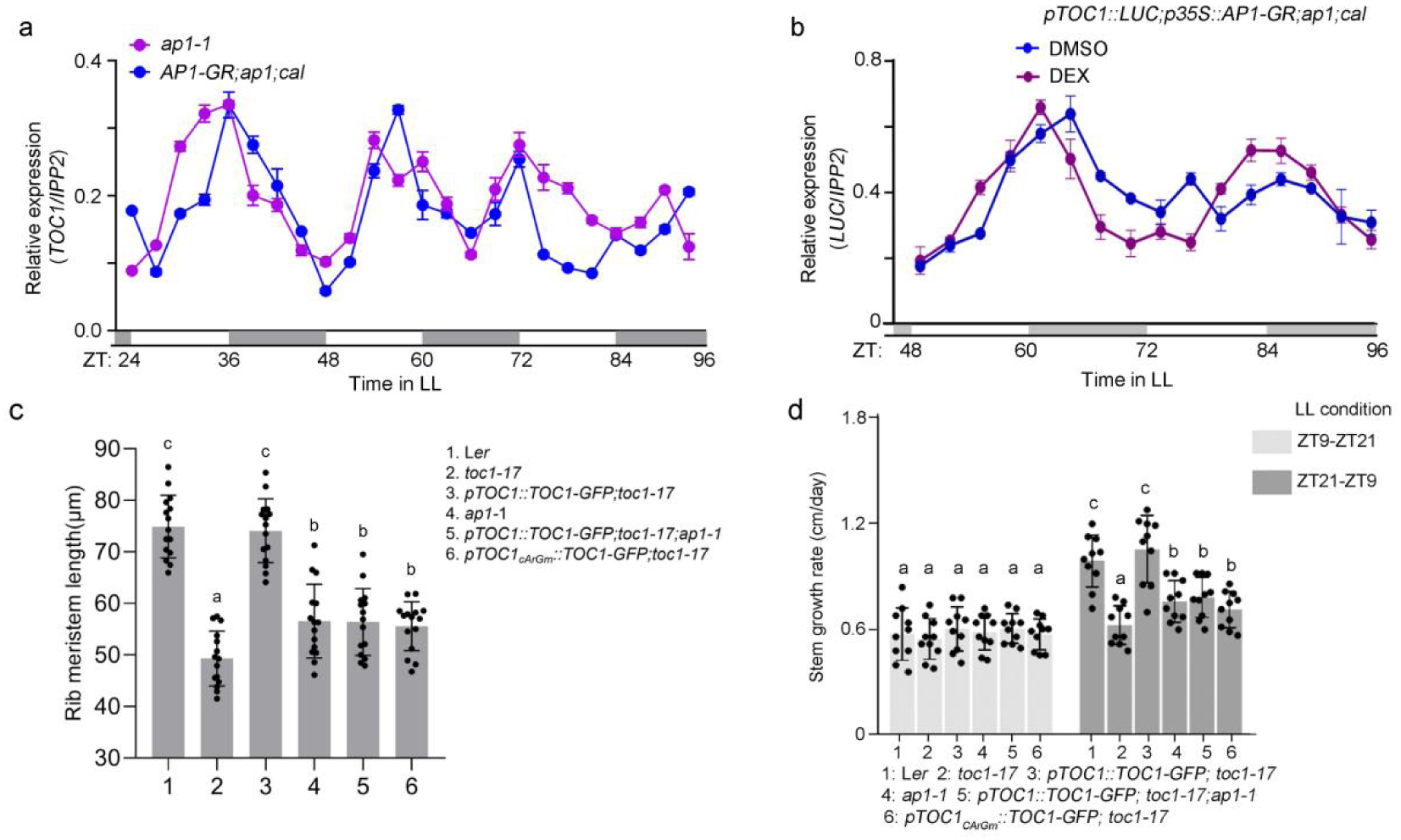
AP1 control *TOC1* expression. (**a** and **b**) RT-qPCR to examine the circadian expression pattern of *TOC1* in the meristems of indicated plants (**a**) and the *LUC* expression in *pTOC1::LUC;AP1-GR;ap1;cal* under DMSO or DEX treatments (**b**). Meristem sampling was conducted as shown in Fig. 3a at 3-hour interval. Data represent mean ± SD. of three biological replicates. *IPP2* was used as a normalization control. White and gray bars represent subjective day and night, respectively. (**c** and **d**) RM length (**c**) and stem growth rate (**d**) of plants shown in Fig.4g. The letters indicated the significance groups at P < 0.05 (one-way ANOVA and Tukey test). Different letters between genotypes and treatments indicate significance from samples (n=15) in (**c**) and (**d**).

**Extended Data Fig 4.**
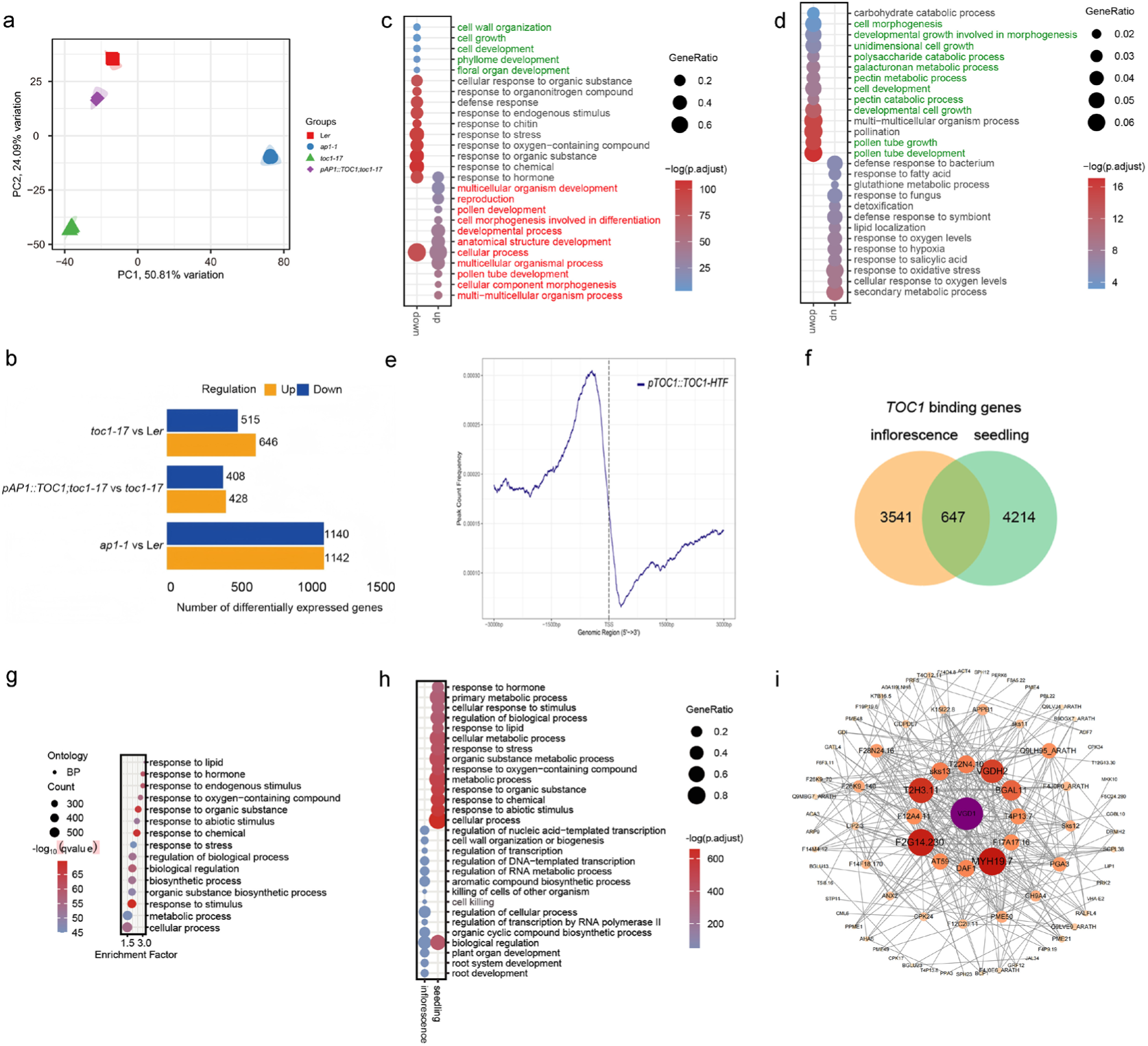
Transcriptomic and ChIP analyses related to. Fig. 1 **and** Fig. 5. (**a**) Principal component analysis (PCA) showing the distribution of variation among samples. PC1 and PC2 account for 50.81% and 24.09% of the total variation, respectively. (**b**) Number of differentially expressed genes (upregulated and downregulated) in each comparison group: *toc1-17* vs. L*er*; *pAP1::TOC1;toc1-17* vs. *toc1-17*; *ap1-1* vs. L*er*. (**c** and **d**) Dot plot showing GO enrichment results of upregulated genes and downregulated genes (**b**) in *ap1-1* vs. L*er* (**c**) and *toc1-17* vs. L*er* (**d**). (**e**) Density plot of putative TOC1 binding peaks within ± 3kb of the Transcription start sites (TSS). (**f**) TOC1-binding sites were identified in both seedling and inflorescence samples. (**g**) Dot plot showing GO enrichment results of 647 shared TOC1-binding sites were identified in both seedling and inflorescence samples. (**h**) Dot plot showing GO enrichment results of 3,541 and 4,214 sites were uniquely bound by TOC1 specifically in inflorescences and seedlings. (**i**) A protein-protein interaction (PPI) network analysis on all DEGs identified in *toc1-17* VS. L*er*. Genes with established interactions were retained for network visualization, and *VGD1*, a direct target gene of TOC1 identified by ChIP, was annotated. Our analysis revealed that the interaction network was primarily composed of the PME family and other cell wall-related gene families.

**Extended Data Fig.5.**
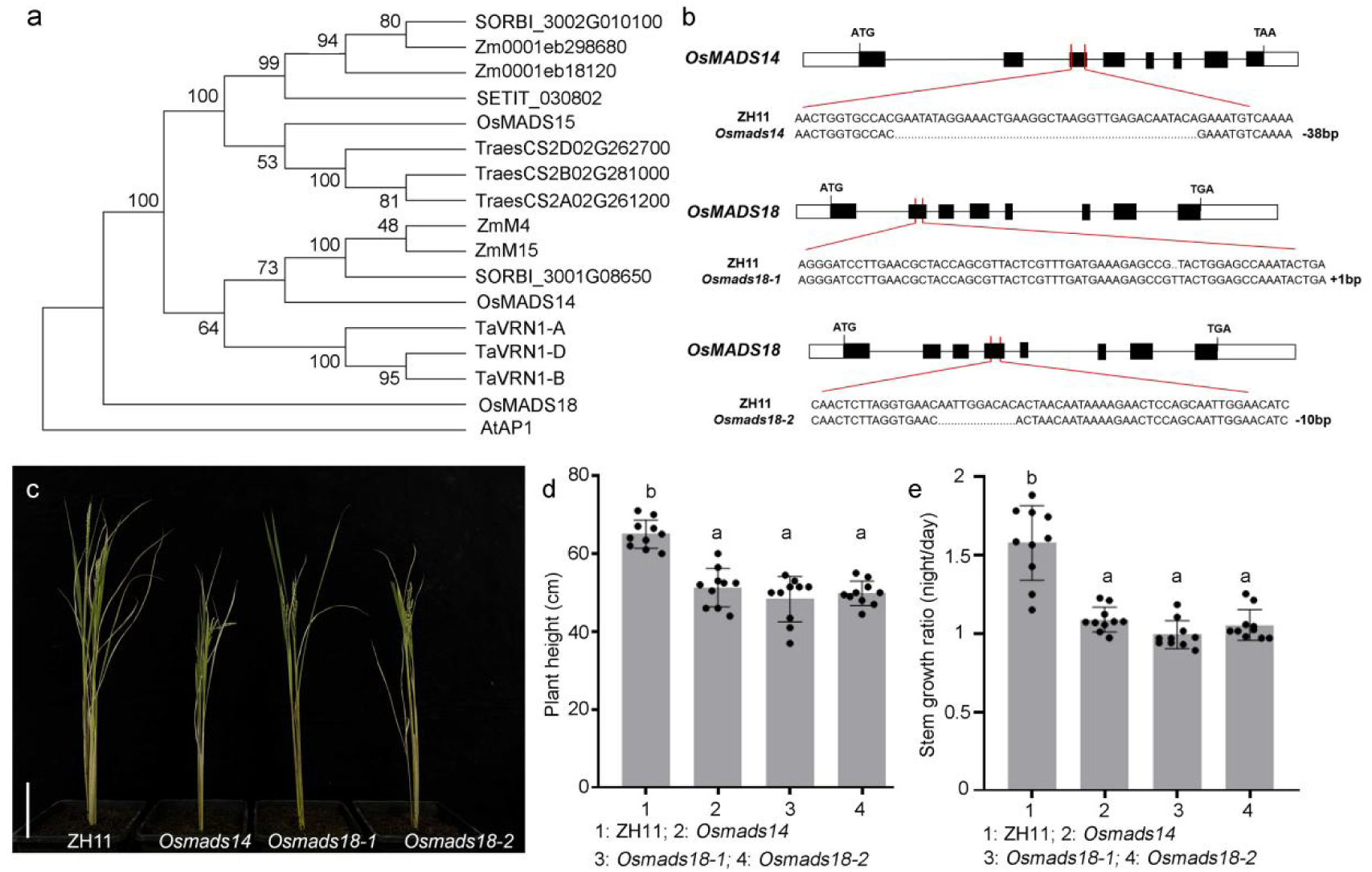
AP1 homologs regulate stem development in rice. (**a**) Phylogenetic tree analysis of AP1 homologous genes. (**b**) Information on the gene editing of rice *MADS14* and *MADS18*. (**c**) Morphological observation of rice ZH11 (WT), *mads14*, *mads18-1*, and *mads18-2*. Scale bars: 10 cm. (**d** and **e**) The peduncle length (**d**) and stem growth ratio (**e**) of rice accessions in (**c**). Data represent mean ±SD (n=10). Different letters indicate statistically significant differences by one-way ANOVA (Tukey’s multiple comparisons test, P < 0.01.

